# CREEPING STEM1 Regulates Directional Auxin Transport for Lodging Resistance in Soybean

**DOI:** 10.1101/2024.06.24.600431

**Authors:** Zhiyong Xu, Liya Zhang, Keke Kong, Jiejie Kong, Ronghuan Ji, Jun Liu, Hongyu Li, Yulong Ren, Wenbin Zhou, Tao Zhao, Tuanjie Zhao, Bin Liu

**Affiliations:** Key Laboratory of Biology and Genetics Improvement of Soybean, Ministry of Agriculture / Zhongshan Biological Breeding Laboratory (ZSBBL) / National Innovation Platform for Soybean Breeding and Industry-Education Integration / State Key Laboratory of Crop Genetics & Germplasm Enhancement and Utilization / College of Agriculture, Nanjing Agricultural University, Nanjing 210095, China; State Key Laboratory of Crop Gene Resources and Breeding, Institute of Crop Science, Chinese Academy of Agricultural Sciences, Beijing, China; Beijing Dabeinong Technology Group Co., Ltd

## Abstract

Soybean, a staple crop on a global scale, frequently encounters challenges due to lodging under high planting densities, which results in significant yield losses. Despite extensive research, the fundamental genetic mechanisms governing lodging resistance in soybeans remain elusive. In this study, we identify and characterize the *Creeping Stem 1* (*CS1*) gene, which plays a crucial role in conferring lodging resistance in soybeans. The *CS1* gene encodes a HEAT-repeat protein that modulates hypocotyl gravitropism by regulating amyloplast sedimentation. Functional analysis reveals that the loss of *CS1* activity disrupts polar auxin transport, vascular bundle development, and the biosynthesis of cellulose and lignin, ultimately leading to premature lodging and aberrant root development. Conversely, increasing *CS1* expression significantly enhances lodging resistance and improves yield under conditions of high planting density. Our findings shed light on the genetic mechanisms that underlie lodging resistance in soybeans and highlight the potential of *CS1* as a valuable target for genetic engineering to improve crop lodging resistance and yield.

## INTRODUCTION

Soybean (Glycine max) is a major source of plant protein and oil, with significant economic value worldwide. Yield increases can be achieved by increasing planting densities, but high densities can lead to lodging, which severely restrict yield per unit area (Liu et al., 2020). Lodging is classified into different levels based on the angle of the stem from the vertical direction, with complete lodging occurring when the stem lies flat on the ground (Pinthus, 1974; Orf et al., 1999). Lodging can occur at different stages of growth and has varying effects on yield (Noor and Caviness, 1980; Baker et al., 2014). Despite some studies investigating soybean lodging traits (Antwi-Boasiako, 2017; Chen et al., 2017; Hwang and Lee, 2019), there have been few reports on the cloning of genes regulating these traits and their potential application in soybean breeding.

Gravity is a dominant external factor that affects plant upright growth (Gilroy, 2008; Wyatt and Kiss, 2013). Deficiencies in perception of gravity can result in varying degrees of tilting (Muthert et al., 2020), and studies have revealed that gravity response process in plants involves four stages: gravity perception, signal transduction, auxin distribution, and organ bending (Hashiguchi et al., 2013; Vandenbrink and Kiss, 2019). The perception of gravity is achieved through specific plant tissues and cells, such as the columella cells in the root system and the endodermis cells in the flower stem and hypocotyl (Su et al., 2017; Nakamura et al., 2019a). Vacuoles in these cells contain amyloplasts that play a crucial role in the perception of gravity through sedimentation during the gravity response process (Band et al., 2012; Baldwin et al., 2013; Ge and Chen, 2019). Under gravitational stimulation, the amyloplasts move in the direction of gravity, resulting in changes in vacuole shape and structure (Nakamura et al., 2019b; Zhang et al., 2019), that affect auxin carrier distribution and auxin transport, leading to unequal growth and bending of the gravity-responsive tissue (Han et al., 2020; Zhu et al., 2020).

The SGRs gene family is primarily responsible for the gravity response in Arabidopsis (Yamauchi et al., 1997; Fukaki and Tasaka, 1999; Zheng et al., 1999; Kato et al., 2002; Yano et al., 2003; Morita et al., 2006; Hashiguchi et al., 2014). Mutations in *SGR1* and *SGR7* result in defects in gravity perception in the endodermis cells and the disappearance of amyloplasts, leading to a loss of gravity sensitivity in the hypocotyl and inflorescence stem. Other mutations in *SGR2*, *SGR3*, *SGR4*, *SGR5*, *SGR6* and *SGR8* can cause abnormal amyloplast accumulation or distribution, leading to abnormal bending of the inflorescence stem or hypocotyl (Kato et al., 2002; Yano et al., 2003; Morita et al., 2006; Hashiguchi et al., 2014).

The unequal distribution of auxin during gravitropism formation is mainly caused by the altered polar auxin transport (PAT) process between cells, which is mediated by the plasmodesmal auxin influx carrier and auxin efflux carrier proteins. These carriers are responsible for the moving of auxin in vasculature from stem to root tip, and then from root tip to both sides of root base (Muday and DeLong, 2001; Petrasek and Friml, 2009; Naramoto, 2017). In this process, the auxin influx carrier AUX1/LAX (Swarup and Bhosale, 2019), auxin efflux carrier PIN (Sauer and Kleine-Vehn, 2019; Teale et al., 2021), and the bidirectional auxin transport carrier ABCB/PGP (Geisler et al., 2017), are responsible for auxin transport. The asymmetrical distribution of PIN proteins in the cell membrane is a crucial factor in determining the direction of auxin transport (Wisniewska et al., 2006). The establishment of auxin concentration gradients is important for the diversification of auxin signaling and its role in organogenesis, vasculature formation, and phototropism (Teale and Palme, 2018; Skokan et al., 2019; Mäkilä et al., 2023).

In this study, we aimed to identify and characterize a gene, named *Creeping Stem 1* (*CS1*), crucial for lodging resistance in soybeans. Through map-based cloning, we identified the *CS1* gene and found that its disruption led to premature lodging of the plant, specifically at the base of the stem in the hypocotyl region. We also observed that the hypocotyl gravitropism was impaired and the rate of starch sedimentation in the inner layer of the endosperm was reduced in the cs1 mutant.

Further investigation revealed that CS1 plays a significant role in the directional transport of auxin, which influences plant morphogenesis. Overexpression of the *CS1* gene resulted in enhanced resistance to lodging and improved yield under high-density planting conditions. Overall, our findings provide new insights into the function of *CS1* in soybean lodging resistance and highlight its potential for use in crop improvement.

## RESULTS

### Identification of the *cs1* Mutant

To identify genes essential for lodging resistance, we screened an EMS-induced mutagenesis library of the NN1138-2 accession. We identified the *cs1* mutant, which exhibited a sprawling phenotype characterized by slender and soft hypocotyls at the base of the main stem during the vegetative growth stage, contrasting with the NN1138-2 accession (Fig. 1A, Supplementary Fig. 1). Upon reaching maturity, the *cs1* mutant displayed significant reductions in various growth parameters, including main stem length, node number, pod number, seed number, grain weight, and 100-grain weight (Supplementary Fig. 2).

**Figure 1.**
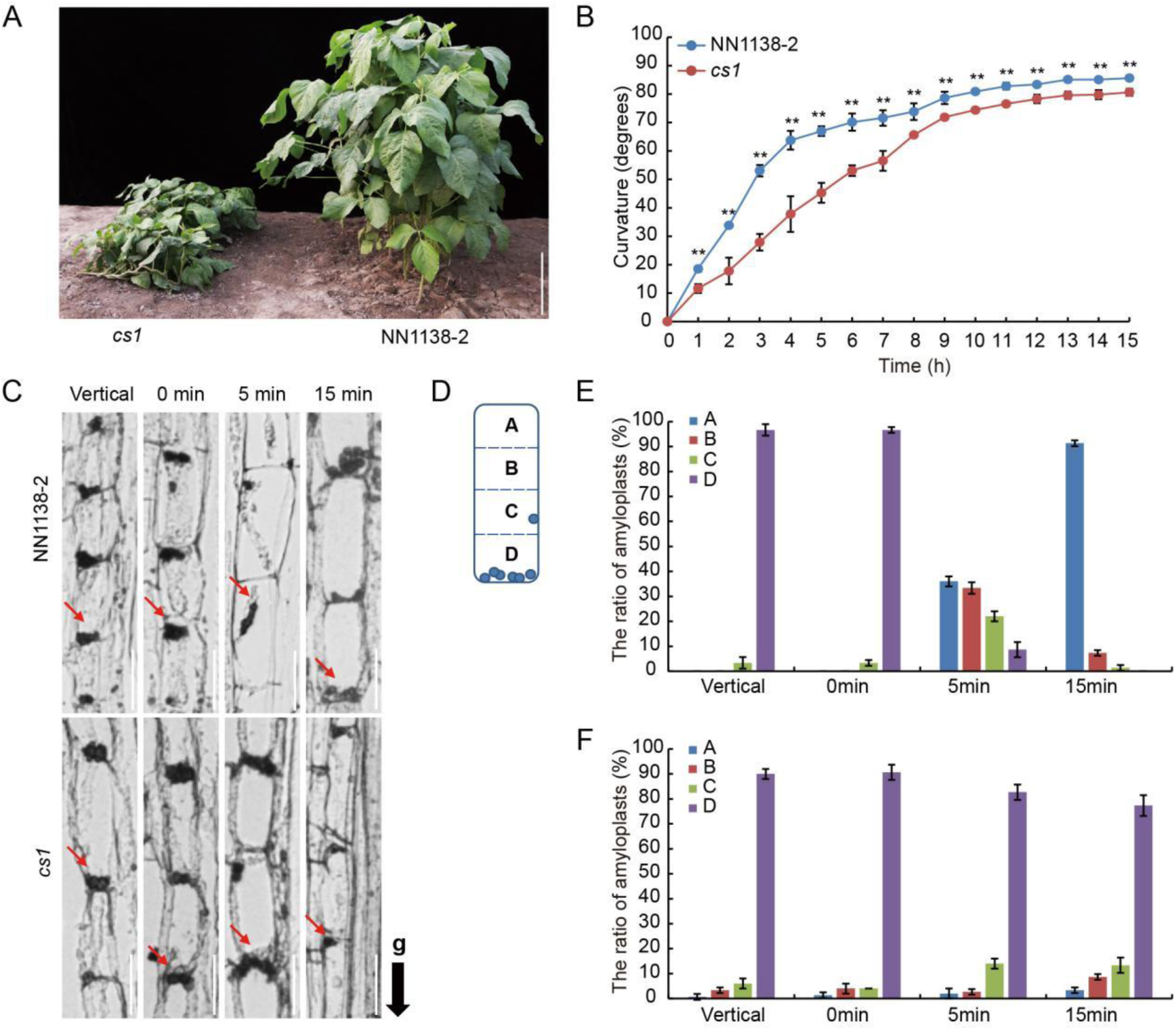
Identification of the *cs1* mutant. **A)** 7-week-old plants of the *cs1* mutant and the wild-type accession NN1138-2 at vegetative stage in the field. Scale bars = 20 cm. **B)** Gravitropic response of hypocotyl in dark. The indicated lines were grown under LD conditions for 7 days. Values are means ± s.d. (*n* = 3). The significant difference between the wild type and *cs1* mutant at each time point was determined by a two-sided *t*-test (** *P* < 0.01). **C)** Longitudinal section images show the distribution of amyloplasts in endodermal cells. The 7-day-old vertical growing plants were inverted for 0 to 15 min. The fragments of hypocotyls (1-2 cm below the cotyledon node) were collected and fixed with the gravity direction maintained at the indicated time point. The red arrows indicate the locations of the amyloplasts, and the black arrow indicates the direction of gravity (g). Scale bars = 5 µm. **D)** Schematic diagram of an endodermal cell, partitioned into four blocks (A, B, C and D from top to bottom) for quantitative analysis, blue circles represent amyloplasts. **E-F)** The ratio of amyloplasts in each block in NN1138-2 (E) and *cs1* (F), Data are means ± s.d. (*n* = 50).

### The *cs1* Mutant Displays Impaired Gravity Response

One of the primary causes of plant lodging is a defect in the gravity response, prompting us to examine the gravity response capability of the *cs1* mutant. We assessed this by subjecting vertically grown seedlings to gravitational stimulation through horizontal placement. Our results showed that the *cs1* hypocotyls demonstrated a reduced bending ability when compared to the wild-type NN1138-2, particularly under dark conditions, with the disparity in bending angle becoming more evident after four hours of stimulation (Fig. 1B, Supplementary Fig. 3). Under light conditions, the bending angles of the *cs1* mutant and wild-type seedlings were only slightly different during the initial three hours, after which they converged (Supplementary Fig. 4). These findings indicate that the *cs1* mutant’s hypocotyl is specifically compromised in gravitropism, with no significant impact on phototropism.

### Amyloplast Sedimentation in the Hypocotyl is Impaired in the *cs1* Mutant

The sedimentation of amyloplasts in the endodermis is crucial for the gravitropic response in plants (Band et al., 2012; Baldwin et al., 2013). To determine the cause of the gravitropic defect in the cs1 mutant, we examined the distribution and sedimentation of amyloplasts within the endodermal cells of the hypocotyl. Bending occurs in the elongation zone of the hypocotyl in response to gravity, prompting us to focus our investigation on amyloplast sedimentation in this area (Supplementary Fig. 5). We inverted the hypocotyl from its normal vertical growth position by 180° and allowed it to incubate for 15 minutes. Subsequently, we employed histological techniques to assess the sedimentation of amyloplasts in the endodermis. For precise evaluation, we divided each endodermal cell into four equal regions, labeled from top to bottom as regions A, B, C, and D (Fig. 1D). In the normal vertical position, we observed that nearly all amyloplasts in both the wild-type NN1138-2 and the *cs1* mutant were situated at the lower part of the endodermal cells (Fig. 1C). Immediately after inversion (0 minute), the amyloplasts in both the wild-type NN1138-2 and the *cs1* mutant were predominantly found at the bottom of the endodermal cells, that is, in region D (Fig. 1C-F). However, as inversion time progressed, the amyloplasts in the wild-type NN1138-2 gradually settled from region D to regions C, B, and A within 5 minutes, whereas most amyloplasts in the *cs1* mutant remained in region D (Fig. 1C, E, F). After 15 minutes of inversion, 91% of the amyloplasts in the wild-type NN1138-2 had moved from region D to region A, whereas 77% of the amyloplasts in the *cs1* mutant remained concentrated in region D (Fig. 1C, E, F). These findings indicate a marked reduction in the sedimentation velocity of amyloplasts in the *cs1* mutant, suggesting that the *CS1* gene facilitates the sedimentation of amyloplasts in the direction of gravity.

### Genetic Analysis of the *cs1* Mutant

To elucidate the genetic basis of the *cs1* mutant’s phenotype, we conducted crosses with four distinct varieties: Williams82, NG94-156, KF1, and NN1138-2, resulting in the creation of four genetic populations. Analysis based on Mendelian genetics revealed that the F_1_ progeny from all four crosses displayed a normal erect phenotype. However, the F_2_ generation exhibited trait segregation that did not adhere to the anticipated 3:1 ratio (Supplementary Table 1). Notably, the F_2:3_ family rows from the crosses of NN1138-2 × *cs1* and KF1 × *cs1* showed a 2:1 ratio of segregated to non-segregated rows (χ² = 0.02, P = 0.88; χ² = 0.01, P = 0.94, Supplementary Table 1). This finding suggests that the prostrate phenotype associated with the *cs1* mutant is likely governed by a single pair of recessive alleles.

### Fine Mapping of the *CS1* Locus

To pinpoint the genes responsible for the prostrate phenotype observed in the *cs1* mutant, we undertook a fine mapping study utilizing the Williams 82 × *cs1* F_2_ population and 1,015 SSR molecular markers. Employing bulked segregant analysis (BSA) (Zhao et al., 2017), we identified three SSR markers, Sat561 (BARCSOYSSR_19_1230), Sat_099 (BARCSOYSSR_19_1251), and Sat_286 (BARCSOYSSR_19_1329), that were tightly linked to the *CS1* locus on chromosome 19 (Supplementary Table 2). These preliminary mapping results were subsequently validated in the F_2_ populations derived from KF1 × *cs1* and NG94-156 × *cs1* crosses.

For the fine mapping of the *CS1* locus, we selected 20 SSR markers within the region and genotyped 42, 89, and 212 prostrate plants from the respective F_2_ populations of Williams 82 × cs1. Analyzing the genotyping data enabled us to delimit the cs1 locus to a 187 Kb interval flanked by two SSR markers, S1290 (BARCSOYSSR_19_1290) and S1302 (BARCSOYSSR_19_1302) (Fig. 2A). Among the 22 candidate genes within this interval (Supplementary Table 3), a single gene, *Glyma.19G186900*, was identified with a nonsense mutation from C to T in the 39th exon of the mutant (Fig. 2A). By sequencing 12 randomly chosen prostrate single plants from the F_2_ population, we confirmed that the base changes were congruent with those observed in the *cs1* mutant (Supplementary Fig. 6). This gene encodes a protein comprising 1,710 amino acids, including six HEAT (Huntingtin, Elongation factor 3, A-subunit of protein phosphatase 2A, and TOR1) repeat units (Supplementary Fig. 7A). Phylogenetic analysis indicated that the homologous gene of *CS1* in Arabidopsis is *SGR6* (*SHOOT GRAVITROPISM 6*), a gene related to gravity sensing (Supplementary Fig. 7B) (Hashiguchi et al., 2014). Quantitative PCR analysis revealed that *CS1* is expressed in all tissues, with the highest expression levels observed in stems (Supplementary Fig. 8). Collectively, our findings posit *Glyma.19G186900* as the candidate gene for *CS1*.

**Figure 2.**
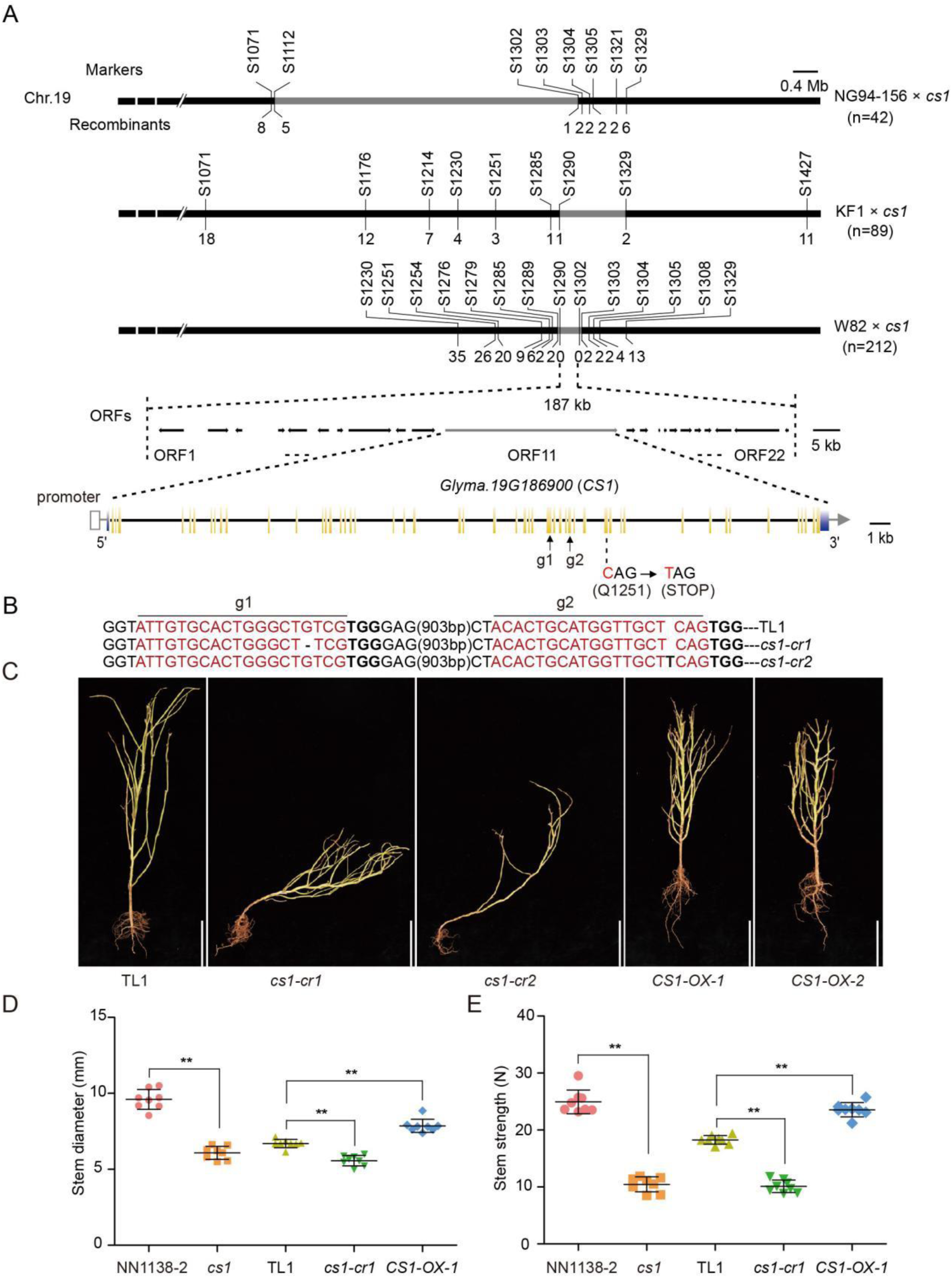
*CS1* encodes a MAESTRO-RELATED HEAT DOMAIN-CONTAINING protein. **A)** Fine mapping of *CS1* candidate gene using three populations: NG94-156 × *cs1*, KF1 × *cs1*, and W82 × *cs1*. The *CS1* locus was narrowed down to a 187-kb region containing 22 annotated ORFs (open reading frame). A nonsense mutation (C to T) was identified in the 39th exon of *Glyma.19G186900* in the *cs1* mutant, leading to a premature protein at Q1252. The *Glyma.19G186900* gene structure is schematically shown; exons and introns are indicated by yellow bars and black bars respectively. The blue bars indicated the 5′- and 3′-untranslated region. **B)** Two sgRNAs (g1 and g2, black arrows) were designed to target the 32th and 36th exons of *Glyma.19G186900* respectively. The mutant sequences of two representative homozygous mutants (*cs1-cr1* and *cs1-cr2*) at T_2_ generation are shown. The target sites of sgRNA are highlighted in red letters with the protospacer-adjacent motif (PAM) in bold. The black base and dash line within the target sites denote nucleotide insertion and deletion respectively. **C)** Plant architectures of the wild type (TL1), *CS1* knockout mutants (*cs1-cr1*, *cs1-cr2*), and *CS1* overexpression lines (*CS1-OX-1* and *CS1-OX-2*) at mature stage in the field. Scale bar = 25 cm. **D-E)** Comparison of hypocotyl diameter (D) and stem length (E) of indicated lines at 46 days after sowing. Data are means ± s.d. (*n* = 8). The significant difference between the indicated line and wild type was determined by two-sided *t*-test (* *P* < 0.05; ** *P* < 0.01).

### *CS1* Gene Dysfunction Leads to Stem Lodging

To ascertain the role of the *CS1* gene, we employed CRISPR/Cas9 gene editing to create two independent homozygous knockout mutants, designated as *cs1-cr1* and *cs1-cr2*. These mutants featured a deletion of the G base at the target site g1 and an insertion of a T base at target site g2, respectively (Fig. 2B). Both mutations led to premature termination of protein synthesis. Field trials revealed that both edited mutants displayed a phenotype identical to that of the *cs1* mutant, characterized by prostrate growth at the stem’s base prior to flowering (Fig. 2C). Furthermore, we generated two *CS1* overexpression lines, *CS1-OX-1* and *CS1-OX-2*, both of which exhibited a dwarf phenotype (Fig. 2C; Supplementary Figs. 9-10). Upon measuring the stem thickness and strength of representative lines from each generation 46 days post-sowing, it was observed that compared to the wild-type controls NN1138-2 and TL1, the *cs1* and *cs1-cr1* mutants had thinner stems and diminished strength. Conversely, the *CS1-OX-1* overexpression line demonstrated significantly increased stem thickness and strength, suggesting enhanced resistance to lodging (Fig. 2D-E). Amyloplast precipitation assays indicated that *cs1-cr1* behaved similarly to the *cs1* mutant, with a marked reduction in the rate of Amyloplast precipitation. In contrast, the *CS1-OX-1* overexpression line showed an accelerated precipitation rate (Supplementary Fig. 11).

### CS1 Influences Vascular Bundle Cell Structure at the Stem Base

To elucidate the regulatory function of the *CS1* gene at the stem base, we examined the cellular composition of hypocotyl cross-sections. The hypocotyl was stratified into three equal segments from apex to base, designated as the upper hypocotyl (UH), middle hypocotyl (MH), and lower hypocotyl (LH) (Fig. 3A). Paraffin section histology revealed that, in comparison to the wild-type TL1, the *cs1-cr1* mutant displayed a reduction in both the area and the number of cell layers within the xylem. Conversely, there was an expansion in the area and an increase in the number of cell layers in the phloem. Additionally, a decrease in the ratio of xylem to phloem was observed in terms of both area and cell layers across all three hypocotyl segments. In stark contrast, the *CS1-OX-1* overexpression line exhibited an enlargement in xylem area without a corresponding increase in the number of xylem cell layers. There was also no significant alteration in phloem area or cell layer count. The xylem to phloem area ratio increased, whereas the ratio of xylem to phloem cell layers remained unchanged (Fig. 3B-H; Supplementary Fig. 12 A-G). Collectively, these findings suggest that *CS1* plays a pivotal role in modulating the development of xylem and phloem within the vascular bundle.

**Figure 3.**
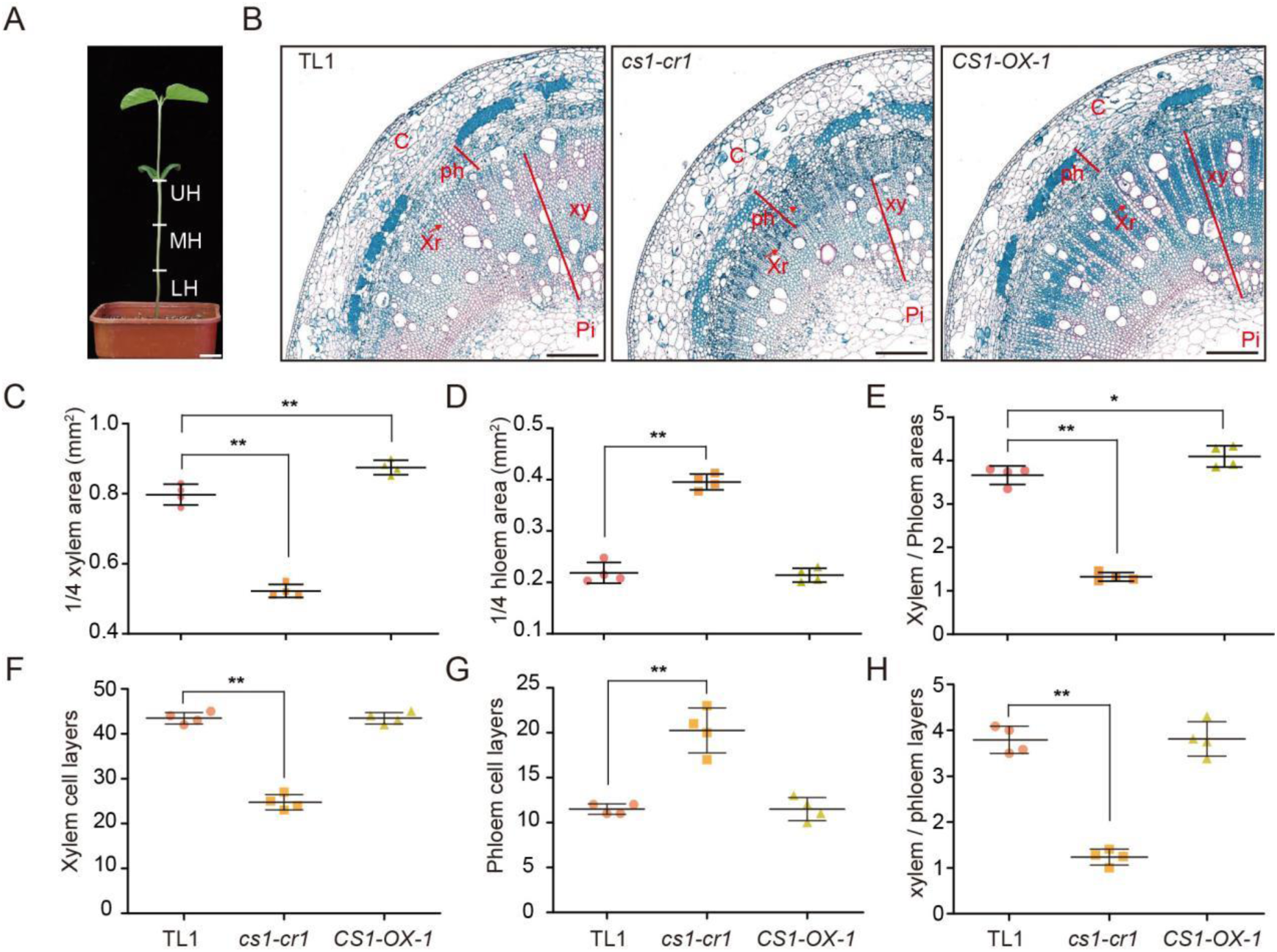
The *CS1* gene regulates xylem and phloem development in hypocotyl. **A)** Pattern of hypocotyl Segmentation. **B)** Quadrant cross section images show the cell layer structures in the upper region of hypocotyls. The plants of indicated lines were grown under LD conditions for 7 days. C, cortex; ph, phloem; xr, xylem rays; xy, xylem; pi, pith. The red arrows represent xylem rays. Scale bar = 250 µm. **C-H)** Scatter plots of xylem areas (C), phloem areas (D), xylem areas / phloem areas (E), xylem cell lays (F), phloem cell lays (G), and xylem cell lays / phloem cell lays (H) of indicated lines as in (B). Data are means ± s.d. (*n* = 4). The significant difference between the indicated line and wild type was determined by two-sided *t*-test (\**P* < 0.05; \*\**P* < 0.01).

### CS1 Influences Cellulose and Lignin Composition in the Hypocotyl

To explore the reasons for the changes in hypocotyl tissue structure in the *cs1* mutant, we conducted a comparative analysis of gene expression profiles across different hypocotyl sections (LH, MH, and UH) between the wild-type NN1138-2 and the *cs1* mutant using transcriptome sequencing. An increment gradient in the number of differentially expressed genes was observed from the LH to the UH (Fig. 4A). Specifically, in the UH, the count of upregulated genes was approximately double that observed MH or LH, and the downregulated genes were roughly triple in number (Fig. 4A). This suggests that the absence of *CS1* function exerts a more significant impact on the UH section of the hypocotyl. Subsequent Gene Ontology (GO) pathway analysis of differentially expressed genes in the UH section highlighted an enrichment of cell wall-related genes (Fig. 4B). These included Expansins, which facilitate cell wall expansion (Marowa et al., 2016), Extensins, which are integral to cell wall assembly and growth in cells (Sede et al., 2018), and xyloglucan endotransglucosylase/hydrolase (XTH), which modulate cell wall tension and rigidity (Eklof and Brumer, 2010), (Fig. 4C). A substantial number of genes with these families exhibited differential expression between the wild-type NN1138-2 and the *cs1* mutant across hypocotyl sections, with particularly pronounced upregulation observed in the UH section of the mutant (Fig. 4C, Supplementary Fig. 13). In line with the transcriptomic data, transmission electron microscopy revealed a marked reduction in the secondary cell wall thickness of the endodermal cells in the *cs1* mutant (Supplementary Fig. 14 A, B). Furthermore, there was a significant decrease in both cellulose and lignin content across all hypocotyl sections of the *cs1* mutant (Supplementary Fig. 14 C, E), whereas in the *CS1-OX-1* overexpression line, these components were significantly elevated (Supplementary Fig. 14 D, F).

**Figure 4.**
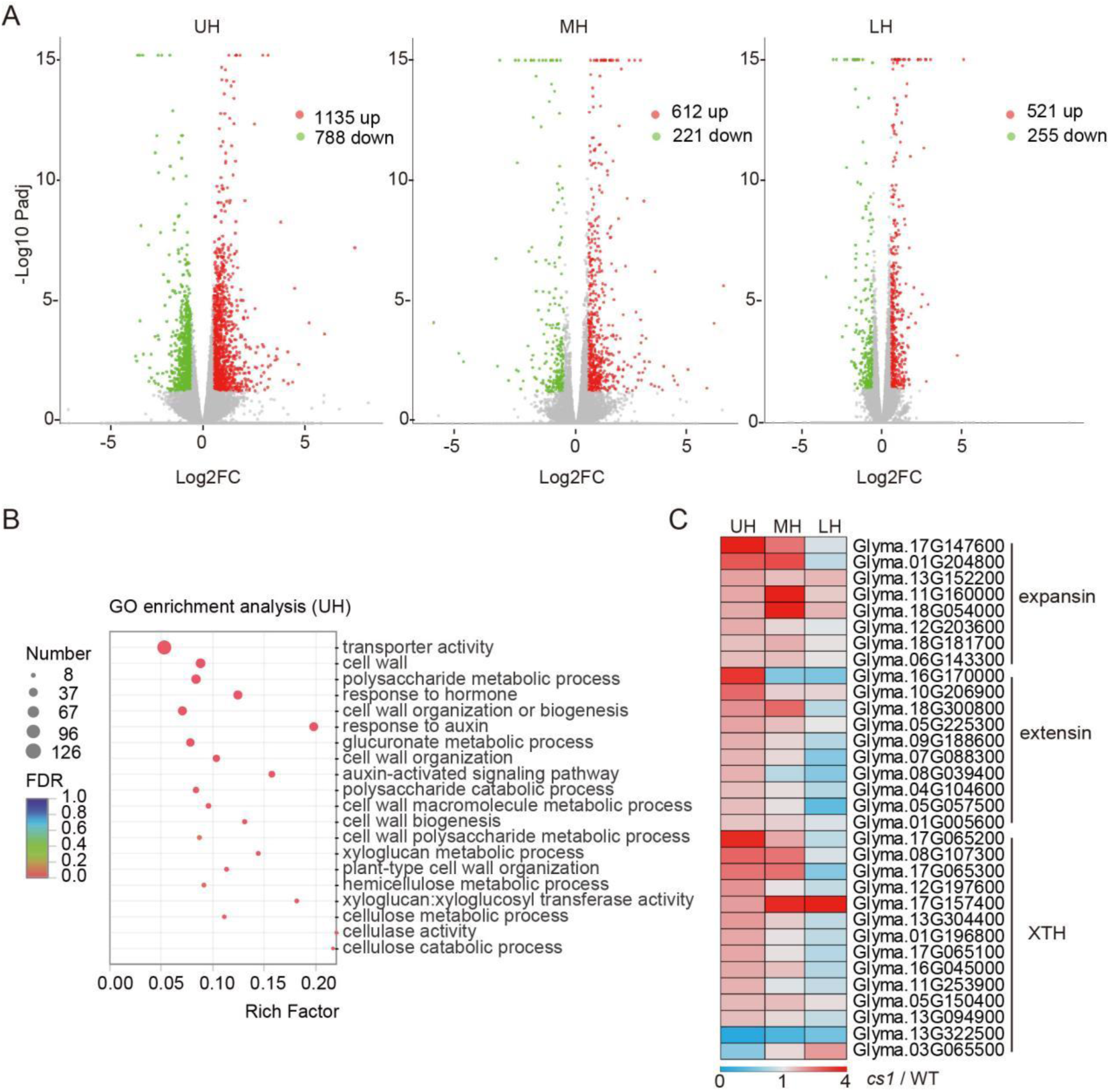
The *CS1* gene affects cell wall related genes and auxin transport related genes changes in the upper hypocotyl. **A)** Volcano plot of gene expression differences between the NIL lines in the indicated segments of hypocotyls. The abscissa is the multiple of the difference of gene / transcript expression between the two samples, and the ordinate is the statistical test value of the difference of gene expression, namely p value. Each dot in the plot represents a specific gene, the red dot represents the significantly up-regulated gene, the blue dot represents the significantly down-regulated gene, and the gray dot represents the non-significant differential gene. **B)** GO term related to cell wall and auxin in UH. The GO terms related to cell wall and auxin in UH were plotted by GO IDs. The vertical axis represents the GO term, horizontal axis represents the Rich factor [the ratio of the number of genes / transcripts enriched in the GO term (Sample number) to the number of annotated genes / transcripts (Background number)]. The larger the Rich factor, the greater the degree of enrichment. The size of the dot indicates the number of genes / transcripts in the GO Term, and the color of the dot corresponds to different Padjust (*P* value-corrected) ranges. **C)** Heat map of expansin, extensin and XTH gene families (involved in cell wall organization and modification) in UH, MH and LH. Plotted with the TPM*^cs1^*/TPM^WT^ of the corresponding part, red represents up-regulation > 1, light blue represents down-regulation < 1, and gray represents no change = 1.

### CS1 Promotes Auxin Concentration Gradient Formation in the Hypocotyl

The GO pathway analysis of transcriptome data also revealed significant enrichment of genes related to auxin pathway. We conducted a detailed examination of genes associated with auxin synthesis (Yue et al., 2014), auxin metabolism (Ljung, 2013), auxin transport (Petrasek and Friml, 2009), and auxin response (Stortenbeker and Bemer, 2019) across different sections of the hypocotyl. The number of differentially expressed genes progressively increased from those involved in auxin synthesis to auxin response. Notably, auxin response genes, including *GmSAURs* and *GmIAAs*, were upregulated in both the UH and MH sections of the *cs1* mutant.

Interestingly, within the upregulated genes in the UH section, 34 were found to be homologous to *GmSAUR14* (Fig. 5A, Supplementary Fig. 15). We randomly selected five genes related to auxin transport and response for qRT-PCR validation, which confirmed the transcriptome sequencing results (Supplementary Fig. 16). We hypothesize that the up-regulation of auxin response genes may be due to an increase in auxin content. To test this hypothesis, we quantified the endogenous auxin levels across each hypocotyl section. The results showed that the auxin levels in both NN1138-2 and TL1 increased progressively from the UH to the LH sections. In contrast, the auxin gradient in the *cs1* and *cs1-cr1* mutants was disrupted, with higher auxin levels observed in each section compared to their wild-type counterparts (Fig. 5B, C). The *CS1-OX-1* overexpression line exhibited an auxin content gradient similar to the wild-type TL1, albeit with significantly reduced levels in each section (Fig. 5B, C). Collectively, these results suggest that CS1 may enhance polar auxin transport, leading to the establishment of an auxin concentration gradient from the apex to the base of the hypocotyl.

**Figure 5.**
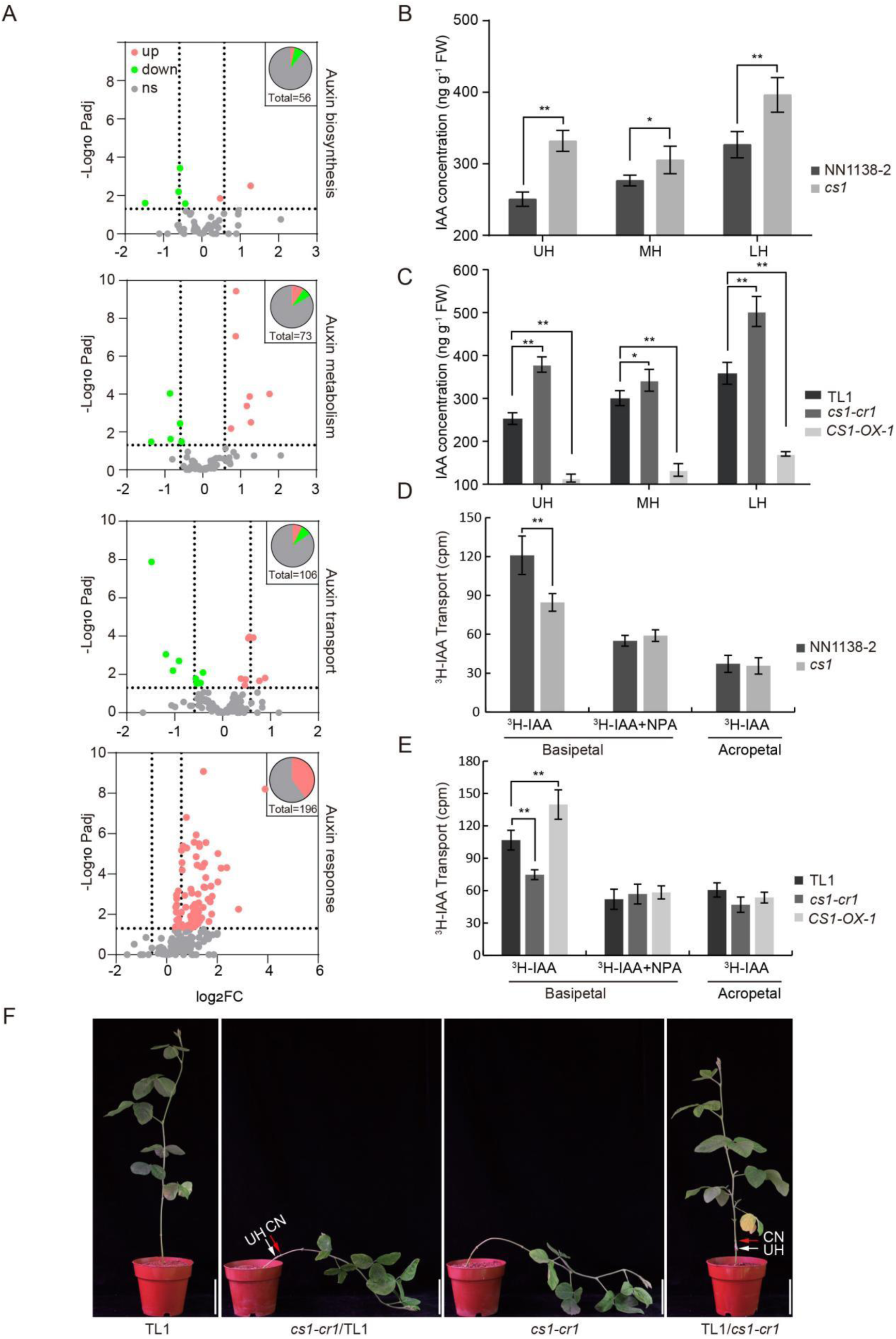
The slow PAT in *cs1* results in the increase of auxin concentration and gradient disorder. **A)** Volcanic map of auxin biosynthesis, metabolism, transport and response related genes in UH. The abscissa represents the fold change of gene expression difference between the two samples, while the ordinate represents the statistical test value (p-value) of the difference in gene expression change. Each point in the figure represents a specific gene, where red dots indicate significantly up-regulated genes, green dots indicate significantly down-regulated genes, and gray dots represent non-significantly different genes. The two dashed lines on the abscissa represent 1.5 times up- and downregulation of genes, respectively, while the dashed line on the ordinate represents p = 0.05. **B-C)** Auxin concentration in each segment of hypocotyls of indicated lines. FW means fresh weight. Values are means ± s.d. (*n* ≥ 5). **D-E)** Comparison of PAT differences in hypocotyls among indicated lines. Values are means ± s.d. (*n* =5). Comparison of differences in PAT among TL1, *cs1-cr1*, and *CS1-OX-1*. Values are means ± s.d. (*n* =5). B-E, The asterisks indicate statistically significant difference from the wild type by two-sided *t*-test (* P <0.05; ** P <0.01). **F)** Grafting leads to *cs1-cr1* not lodging, but TL1 lodging. The white arrow indicates the grafting position, and the red arrow indicates the cotyledon node. CN means cotyledon node. Representative images of at least three independent grafting experiments of indicated combination. *cs1-cr1*/TL1: *cs1-cr1* is scion, TL1 is rootstock; TL1/ *cs1-cr1*:TL1 is scion, *cs1-cr1* is rootstock. Scale bar = 10 cm.

### CS1 Enhances Polar Auxin Transport within the Hypocotyl

To explore the reasons behind changes in auxin content and the disruption of its gradient within the *cs1* mutant hypocotyl, we conducted experiments on polar auxin transport. This involved measuring the rate of polarity transport of IAA (indole-3-acetic acid) labeled with H^3^ from the apical to the basal end of the UH section. The results revealed that the auxin transport rate in the *cs1* mutant was substantially lower compared to the wild-type NN1138-2 (Fig. 5D). A comparable reduction was observed when comparing the edited mutant *cs1-cr1* with the wild-type TL1. In contrast, the auxin transport rate was significantly higher in the *CS1-OX-1* overexpression line (Fig. 5E). These observations indicate that CS1 likely enhances polar auxin transport, which could be a direct cause of the observed changes in auxin content and gradient within the hypocotyl.

### CS1 Modulates Auxin Redistribution under Gravitational Stimuli

To determine whether the insensitivity of the *cs1* mutant to gravitational stimuli was linked to disruptions in polar auxin transport, we conducted an experiment with soybean seedlings exhibiting normal upright growth. The plants were subjected to gravity stimulation by positioning them horizontally, which allowed us to delineate the elongation zone (UH section) of the hypocotyl into lower and upper sides along the longitudinal axis (Supplementary Fig. 17A).

Subsequently, we monitored changes in auxin distribution across these sides over various durations of horizontal positioning. Our findings revealed that with extended periods of gravity stimulation, there was an uneven increase in auxin content on both the lower and upper sides of the wild-type NN1138-2, with a notably higher auxin content on the lower side (Supplementary Fig. 17B). This resulted in asymmetric elongation between the upper and lower sides of the hypocotyl, causing a curvature away from the gravitational pull. In contrast, the *cs1* mutant displayed a similar trend of increasing auxin content within the elongation zone of the hypocotyl. However, the auxin content on the lower side did not significantly exceed that on the upper side (Supplementary Fig. 17C), which may explain the reduced responsiveness of *cs1* to gravitational stimuli.

### CS1 Function in the Upper Hypocotyl Facilitates Lignification in the Lower Hypocotyl

Despite the loss of *CS1* function resulting in the *cs1* mutant’s reduced sensitivity to gravitational stimuli, the hypocotyl of the *cs1* mutant still demonstrates a residual bending response away from the gravitational pull when positioned horizontally. This observation led us to hypothesize that inadequate lignification in the lower hypocotyl may contribute to stem lodging in the *cs1* mutant. Furthermore, we postulated that CS1 might enhance lignification in the lower hypocotyl, either directly or indirectly, through the regulation of auxin transport. To test these hypotheses, we performed grafting experiments by joining the UH sections of 7-day-old wild-type TL1 and the edited mutant *cs1-cr1*. We subsequently observed the tissue structure and quantified the levels of auxin and cellulose in both the scion and rootstock of the grafted plants after a 30-day period.

The results showed that when the wild-type TL1 served as the rootstock and the edited mutant *cs1-cr1* as the scion, the grafted plants exhibited a creeping phenotype, like that of the *cs1-cr1* mutant (Fig. 5F, Supplementary Fig. 18). Additionally, there was a decrease in the xylem-to-phloem area ratio within the vascular bundle of the rootstock below the graft union, accompanied by a reduction in auxin levels and an increase in cellulose content, all of which aligned with the characteristics of the ungrafted *cs1-cr1* mutant (Supplementary Fig. 19). In contrast, when the hypocotyl of the *cs1-cr1* mutant was used as the rootstock and that of the wild-type TL1 as the scion, the grafted plants exhibited a normal upright phenotype resembling the wild-type TL1 (Fig. 5F; Supplementary Fig. 18). Moreover, the xylem-to-phloem area ratio in the rootstock’s vascular bundle increased, and there was a decrease in auxin and an increase in cellulose, all of which mirrored the attributes of the ungrafted wild-type TL1 (Supplementary Fig. 19). These results suggest that the phenotypic changes induced by grafting are likely attributable to modifications in the rootstock’s tissue structure. We hypothesize that the observed alterations in the rootstock’s tissue structure may be mediated by a mobile factor from the scion. Auxin, known for its capacity for translocation and variable concentration levels, is a prime candidate for inducing these structural changes.

### CS1 Modulates Auxin Distribution and Impacts Root Development

To ascertain whether the loss of *CS1* function influences the distribution of endogenous auxin within the roots, we utilized the auxin reporter system *DR5::GUS* to analyze auxin distribution in the roots. Our findings revealed that in comparison to the wild-type control, the *cs1* and *cs1-cr1* mutants displayed reduced GUS expression in both the endodermis and cortex of the roots. Conversely, the *CS1-OX-1* overexpression line exhibited elevated GUS expression levels in these same regions (Fig. 6A). Auxin, a crucial hormone for root development, plays a critical role in root morphogenesis, as changes in its content can significantly affect this process (Du and Scheres, 2018). Examination of root morphology across various stages indicated that at the 7-day-old stage, the *cs1* and *cs1-cr1* mutants presented with shorter primary roots, an increased number of lateral roots, and reduced lengths of the longest lateral roots. In contrast, the *CS1-OX-1* overexpression line demonstrated opposite phenotypic changes (Fig. 6B, D, E, F). By the 50-day-old stage, there was a notable decrease in root biomass for the *cs1* and *cs1-cr1* mutants, whereas the *CS1-OX-1* overexpression line exhibited a more developed root system, characterized by an abundance of newly formed root hairs (Fig. 6C). Collectively, these results imply that CS1 likely influences root morphology and biomass by modulating auxin distribution within the roots.

**Figure 6.**
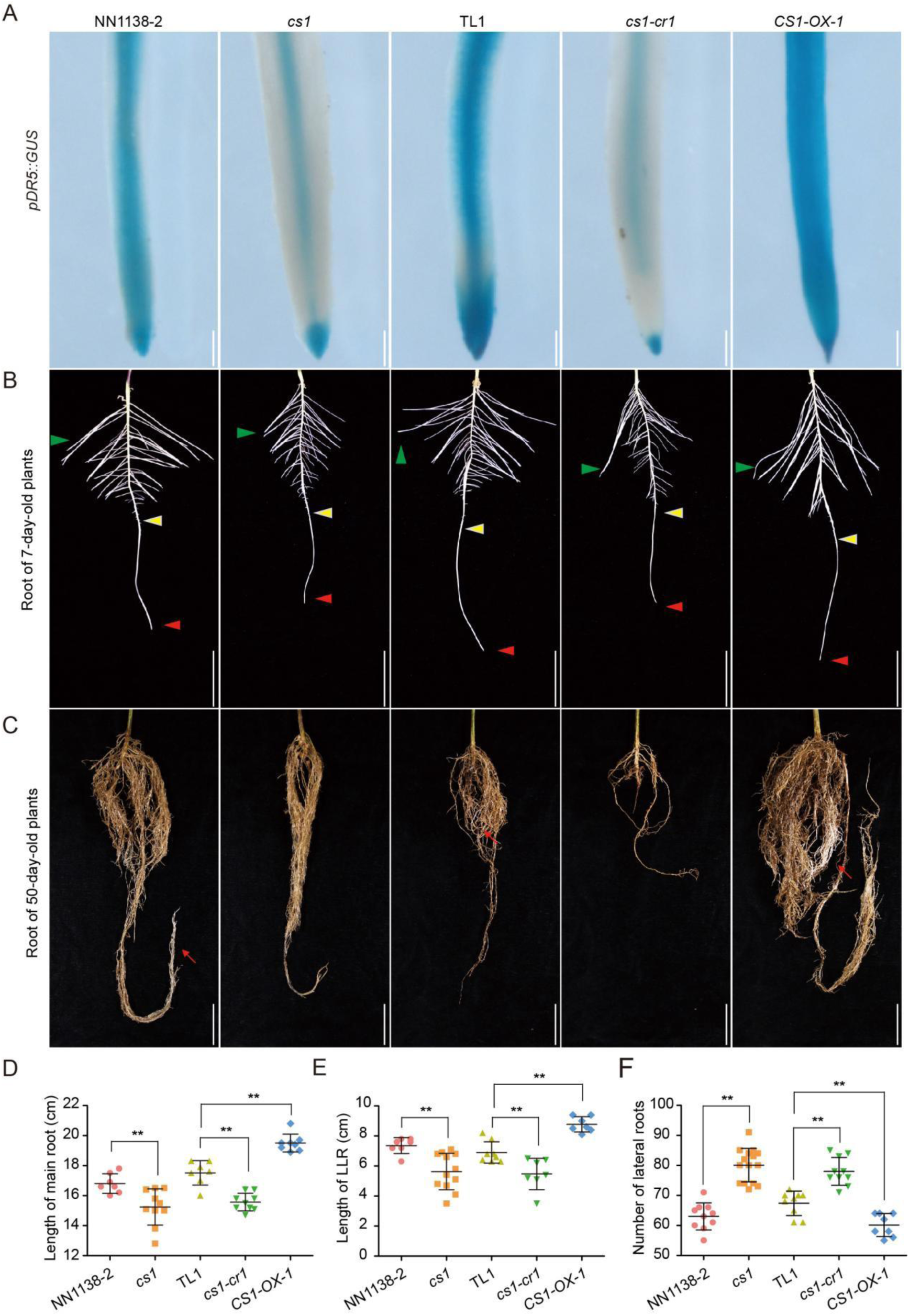
Dysfunction of *CS1* confers abnormal auxin distribution and root morphology. **A)** Histochemical staining of *pDR5::GUS* transgenic hairy roots in the indicated backgrounds. Three independent experiments yielded similar results. Scale bar = 50 µm. **B)** Root images of 7-day-old seedlings grown under LD conditions. The red triangle represents the tip of the main root, the green triangle represents the longest lateral root, and the yellow triangle represents the newest lateral root. Scale bar = 4 cm. **C)** Representative root images of indicated genotypes grown under LD conditions for 50 days. Scale bar = 3 cm. **D-F)** The main root length, the longest lateral root length (data from B), and the number of lateral roots (data from B). Values are means ± s.d. (*n* = 10). The asterisks indicate statistically significant difference from the wild type by two-sided *t*-test (** P <0.01).

### CS1 Overexpression Enhances Soybean Yield under High-Density Planting Conditions

High-density planting is recognized as an effective strategy to boost soybean yields. However, it can lead to reduced blue light penetration, causing plants to elongate and increasing the risk of lodging, which may counteract the intended yield benefits (Lyu et al., 2021). To address this, we assessed the performance of various genotypes under low blue light (LBL) conditions in laboratory settings. Our results indicated that under LBL conditions, both the wild-type TL1 and *cs1* mutant plants experienced lodging around 21 and 25 days post-treatment, respectively. In contrast, the *CS1-OX-1* overexpression line remained erect even beyond the fourth week (Fig. 7A-B). Subsequently, we evaluated the performance of the *CS1-OX* lines under field conditions at planting intervals of 10 cm and 5 cm. The data revealed that the *CS1-OX* lines consistently displayed a phenotype characterized by shorter, thicker stems under both spacing conditions (Fig. 7C). In terms of yield, under the 10 cm interval, the wild-type TL1 and *CS1-OX* lines had comparable yields. However, under the 5 cm interval, the wild-type TL1 exhibited a significantly higher lodging rate compared to the *CS1-OX* lines. Moreover, the yields of the *CS1-OX-1* and *CS1-OX-2* overexpression lines were found to be 15% and 12% higher, respectively, than those of the wild-type TL1 (Fig. 7C). These results suggest that the *CS1* gene holds potential for the development of soybean cultivars that are tailored for higher yield under high-density planting conditions.

**Figure 7.**
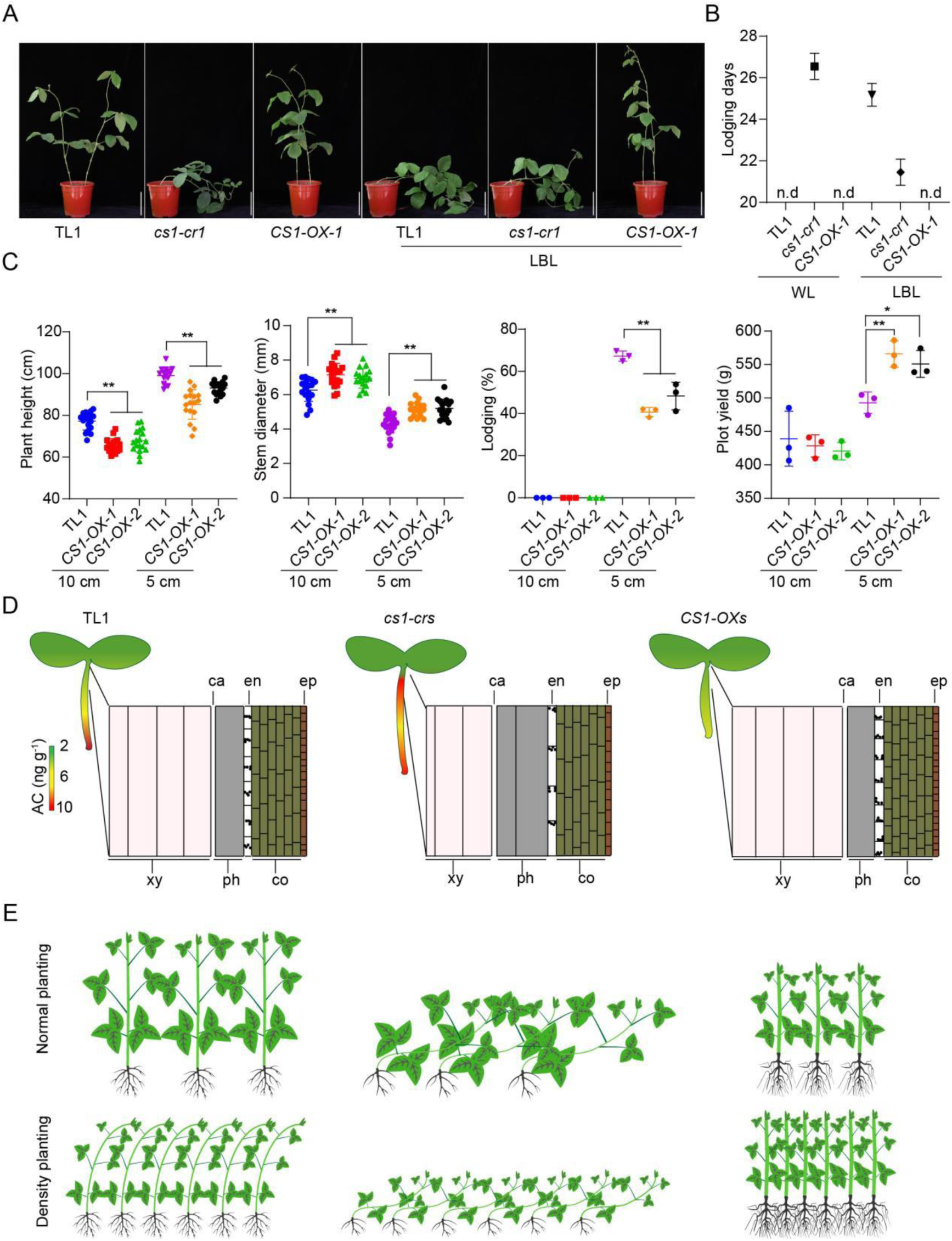
Density planting exacerbated TL1 lodging, but had little effect on *CS1-OXs*. **A)** Lodging phenotype under wight light (WL) and low blue light (LBL). Scale bar = 9 cm. **B)** Plant lodging days in a. Values are means ± s.d. (*n* = 4). **C)** Comparison of the plant height, stem diameter, lodging rate and plot yield of TL1 and *CS1-OXs* under normal planting and density planting. Normal planting means 10 cm space. Density planting means 5 cm space. Plants at maturity stage R8 whose main stem leaned more than 45° were recorded as having lodged. Data are means ± s.d of three biological repeats. The significant difference between the indicated line and wild type was determined by two-sided *t*-test (* P < 0.05, ** P < 0.01). **D)** Auxin concentration gradient and organization structure pattern diagram in TL1, *cs1-cr1* and *CS1-OX-1*. The black circles in en represent amyloplasts, and the position represents the distribution of amyloplasts after plants were inverted 5 minutes. Xy, xylem; ca, the cambium; ph, phloem; en, endodermis; co, cortex; ep, epidermis. AC means auxin concentration. **E)** The morphology of TL1, *cs1-cr1* and *CS1-OX-1* at different planting densities.

## DISCUSSION

Our research explores the genetic complexities that underlie soybean’s resistance to lodging, a trait vital for crop yield and adaptability. Through map-based cloning, we have identified and characterized the *CS1* gene, which is responsible for regulating the creeping stem phenotype in soybean. Functional analysis of the *CS1* gene, using genetic transformation and gene editing techniques, has clarified its pivotal role in shaping plant architecture and enhancing lodging resistance. This discovery provides a molecular target for refining crop improvement strategies.

The *cs1-cr* mutants, generated through targeted gene editing, exhibit a creeping stem phenotype akin to the one observed in the *cs1* mutant generated by EMS mutation. Conversely, CS1 overexpression results in a robust plant architecture with enhanced lodging resistance, particularly beneficial under high-density planting conditions that favor increased yields. Physiological assessments of the *cs1* mutant uncovered disruptions in amyloplast sedimentation within the hypocotyl’s endodermal cells and a diminished gravitropic response. These findings, coupled with a sluggish auxin content adjustment under gravitational stimulus, point to a complex interplay between cellular and molecular mechanisms that govern plant gravitropism. Histological studies further reveal structural perturbations in vascular bundle distribution, which are partially reversible through grafting, suggesting a degree of plasticity in the vascular system’s response to genetic changes.

Transcriptional profiling of the *cs1* mutant has shed light on the molecular mechanisms that underlie the observed phenotypes. Specifically, we observed significant alterations in the expression of auxin transport carriers, including GmPINs and GmABCBs. These transcriptional shifts are hypothesized to result from impaired auxin polar transport, leading to elevated auxin concentrations and disrupted gradients (Fig. 7D, E). Such auxin homeostasis disruptions are likely to impact hypocotyl and root development, contributing to the *cs1* mutant’s increased susceptibility to lodging.

Our research reveals a potential link between the soybean *CS1* gene and the Arabidopsis *SGR6* gene, both integral components of the HEAT-repeat protein family, which play pivotal roles in plant gravity sensing. While the loss of function in both genes attenuates gravitropism, the phenotypic manifestations differ markedly. The *sgr6-1* mutant of Arabidopsis exhibits no significant morphological defects apart from the lateral shoots’ horizontal elongation. In contrast, the absence of *CS1* in soybean leads to a spectrum of developmental alterations, including weakened hypocotyl gravitropism, a slender and soft main stem above ground, impaired root morphology, and a prostrate growth habit before flowering. Both CS1 and SGR6 are implicated in amyloplast sedimentation, a process critical for gravity sensing, suggesting a conserved mechanism in plant adaptation to gravity (Chen et al., 2023; Nishimura et al., 2023). It is plausible that CS1 and SGR6 modulate amyloplast sedimentation, thereby influencing polar auxin transport (PAT) in response to gravitational cues (Supplementary Fig. 20). This hypothesis warrants further investigation to elucidate the intricate mechanisms by which these genes orchestrate plant gravitropic responses.

The discovery of the *CS1* gene enhances our understanding of the molecular underpinnings of lodging resistance in soybean, with far-reaching implications for crop enhancement. Overexpression of *CS1* confers a suite of advantageous agronomic traits, including reduced plant height, increased stem thickness, and enhanced lodging resistance, that render it a prime candidate for genetic engineering initiatives. This gene has the potential to transform the development of soybean cultivars that combine high yield with superior lodging resistance, addressing a significant challenge in soybean cultivation globally. Furthermore, the *cs1* mutant, characterized by its distinct phenotype, provides a unique avenue for exploration of the genetic architecture underpinning lodging resistance and its complex interplay with other economically important traits. The multifaceted role of *CS1* in shaping plant architecture, regulating auxin homeostasis, and its evolutionary conservation across diverse plant species, highlight its fundamental importance in plant biology.

## METHODS

### Plant Materials and Growth conditions

Soybean varieties NanNong1138-2 (NN1138-2), Williams 82 (W82), Kefeng 1 (KF1), NG94-156, and Tianlong 1 (TL1) were obtained from the Nanjing Agricultural University’s Germplasm Resource Bank in Jiangsu Province, China. The *cs1* mutant was derived from EMS mutagenesis of NN1138-2 and bred through high-generation self-breeding. TL1 was used for genetic transformation. Unless otherwise specified, all seedlings were cultivated under long-day (LD) conditions with 16 hours of light and 8 hours of darkness, at a temperature of 25℃, and under white LED lights with an intensity of 90 μmol·m^−2^·s^−1^.

### Gravitropism Assay

Seedlings were grown under LD conditions for 7 days. At the end of the light period, plants were horizontally placed, and hypocotyl curvature was recorded using an intelligent camera (Mijia 1080P, night mode) in darkness. The bending angle between the horizontal baseline and the apex growing direction was measured using the ImageJ program. To determine the gravity sensing region, the hypocotyl of 3.5-day-old seedlings was marked in 5 mm increments from the growth point to the base. After 24 h, each unit was measured, and sections were cut in 5 mm increments from stem apical. The plant was then placed in complete darkness, and the bending state was observed using an intelligent camera for 10 h.

### Amyloplast Sedimentation Assay

Seedlings were grown under LD conditions until cotyledons were fully expanded. The seedlings were then vertically inverted for 0, 5, or 15 min in darkness before fixation in FAA. Paraffin sections were stained with iodine-potassium iodide, following the methods described before (Kim et al., 2011).

### Gene Mapping

Four F_2_ populations were constructed by crossing the *cs1* mutant with NN1138-2, W82, KF1, and NG94-156. A Chi-square test was performed to analyze the segregation ratio in the F_2_ and some F_2:3_ populations. Bulked segregant analysis (BSA) was used to screen polymorphic markers linked to the mutant genes. DNA from 20 normal and 20 mutant plants of the F_2_ population was mixed in equal amounts to create normal and mutant plant gene pools. SSR markers were used to detect polymorphisms between the gene pools, and Mapmaker 3.0 software was used to analyze the F_2_ population for linkage to the target gene. Fine mapping localized the mutation between markers S1289 and S1303 in W82 × *cs1*, S1290 and S1329 in KF1 × *cs1*, and S1112 and S1302 in NG94-156 × *cs1*. Primer sequences for fine-mapping are listed in Supplementary Table 2.

### Phylogenetic Analysis

CS1 homologous proteins from other species were retrieved using a BLASTP tool from NCBI. A neighbor-joining method based on MEGA 7 software was used to construct a phylogenetic tree. Bootstrap analysis with 1000 resamplings was performed to determine branch support.

### Plasmid Construction and Genetic Transformation

To construct the *CS1-YFP* or *YFP- CS1* overexpression vector, the synthetic coding DNA sequence (CDS) of *CS1* (5,130 bp) was inserted into the *pJRH0641* vector by the T4 ligase following the manufacturer’s instructions (Invitrogen). To generate CRISPR/Cas9-engineered *CS1* mutants, sgRNAs were designed using CRISPR-P website (http://crispr.hzau.edu.cn). The 19-bp DNA fragment coding the gRNA was inserted into the *pCas9-AtU6-sgRNA* vector at the *Xbai*1 site. The CRISPR/Cas9 expression vector was constructed with Cas9 CDS driven by *35S* promoter and the designed gRNA transcribed by *Arabidopsis U6* promoter (Tsutsui and Higashiyama, 2017). Above expression plasmids were individually introduced into *Agrobacterium tumefaciens* strain EHA105 and transformed into wild-type soybean (TL1) by implementing the cotyledon-node method (Paz et al., 2006).

### Measurement of Stem diameter and Strength

Stem diameter and strength were quantified by measuring the hypocotyls of the plants 46 days after sowing in the field. Hypocotyl diameter was determined with a digital vernier caliper, while stem strength was assessed using a Japanese AIKOH RZ push-pull meter. The values displayed by the push-pull meter when the hypocotyls were tilted to 45 degrees were recorded. To ensure statistical robustness, a sample size of 8 plants was randomly selected for both measurements.

### Analysis of Hypocotyl Cross-section

For the observation of hypocotyl cross-section, seedlings were grown under LD conditions for 25 days. The morphological upper end was divided into three parts, and the cross-sections were stained with safranin-O-fast green by paraffin sections. The resulting scanned pictures were examined using KFBIO software. The data for xylem and phloem were measured using ImageJ software. Transmission electron microscopy (TEM) analysis was carried out using hypocotyl of 7-day-old seedlings. The UH segment was cross-sectioned by 1 cm and longitudinally cut into two parts, which were then fixed in 3% glutaraldehyde. Material embedding and slicing were conducted, and the established slices were observed and photographed under a JEM-1230 transmission electron microscope.

### Measurement of Auxin and Cellulose Concentrations

To measure the concentrations of auxin and cellulose, seedlings were grown under LD conditions for 7 and 12 days, respectively. The measurement was conducted on five individual plants using the plant auxin (IAA), plant cellulose (CE), and Lignin enzyme-linked immunoassay (ELISA) kit provided by Jiangsu Meimian Industrial Co., Ltd (Chen et al., 2022). To determine the auxin content after gravity stimulation the seven-day-old plants were horizontally placed at the end of light period and incubated for 0, 1, and 4 h in dark, respectively. The curved part of the hypocotyl was divided into upper and lower sides along the midline, and the auxin content of both sides was measured. At least four plants were included in the experiment to ensure accuracy.

### RNA-Seq Analysis

The hypocotyls of seven-day-old NN1138-2 and *cs1* plants were evenly divided into upper (UH), middle (MH), and lower (LH) segments, starting from the upper end to the lower end of their morphology. Three segments of 15 seedlings were collected for total RNA extraction using Trizol (Invitrogen). Subsequently, nine cDNA libraries consisting of three segments, each with three biological replicates, were constructed following Illumina’s standard protocols. These libraries were then sequenced using Illumina Novaseq 6000 (2×150bp read length) by Major Bio (Shanghai, China). RNA-Seq reads were aligned to the soybean reference genome (https://phytozome.jgi.doe.gov/pz/portal.html#!info?alias=Org_Gmax) using TopHat after filtering out low-quality (lowest base score < 20) reads using SeqPrep and Sickle (Trapnell et al., 2009).

The gene expression levels were computed and normalized to TPM (Transcripts Per Million reads). Further, GO functional-enrichment analysis was conducted to determine which DEGs (Differential Expression Genes) were significantly enriched in GO terms at Bonferroni-corrected P-adjust < 0.05, relative to the whole-transcriptome background. The cut-off for significant differential expression was set as P-adjust < 0.05, with up / down-regulated fold-changes of 1.5 or greater. To analyze gene families related to auxin synthesis, metabolism, transport, and response, we conducted extensive searches for Tryptophan metabolism, ABC transporters, UDP-glycosyltransferase, Indole-3-acetate o-methyltransferase, ILR1, PIN, AUX1, GH3, IAA, and SAUR using the cutting-edge Majorbio cloud platform (https://cloud.majorbio.com/). The searched genes underwent rigorous preprocessing to eliminate genes with no expression and significant fluctuations. Subsequently, we extracted the Log_2_FC (*cs1*-UH /NN1138-2-UH) and P-adjust data for the preprocessed genes, and generated a visually appealing volcano map using Graphpad Prism 9.0.0 software.

### qRT-PCR Analysis

For the synthesis of the first complementary DNA and cDNA, DNase-treated total RNA was subjected to reverse transcription using a TRAN reverse transcription kit. Quantitative real-time polymerase chain reaction (qRT-PCR) was conducted on 96-well optical plates, using a SYBR Green RT-PCR kit (Takara) and a Roche Light Cycler 480. The mRNA level of *GmActin11* was utilized as the internal control for normalization. The analysis was performed on three independent biological replicates, with two replicate reactions for each sample.

### Polar Auxin Transport Assay

The assessment of polar auxin transport was conducted with a slight adaptation of a previously established protocol (Yue et al., 2018). Briefly, 2 cm long segments were obtained by cutting hypocotyls from the tip, with 0.2 cm apical ends removed. These segments were pre-incubated in 1/2 MS (pH 5.8) liquid medium for 3 hours to remove endogenous IAA. The apical or basal ends of these segments (for acropetal or basipetal transport assays, respectively) were then immersed in a 0.6-ml Eppendorf tube containing 20 μl of 1/2 MS liquid medium with 200 nM ^3^H-IAA. For the negative control of basipetal transport assays, the specific auxin transport inhibitor N-1-naphthylphtalamic acid (NPA) was introduced at a concentration of 50 nM. The segments were then maintained in the dark at 28℃ for three hours to allow for the inhibitory effect of NPA to manifest. Subsequently, 5-mm sections were extracted from the non-submerged regions of each segment. These sections were transferred into 2 ml of scintillation liquid for 18 hours and then the radioactivity of each section was quantified using a liquid scintillation counter (1450 MicroBeta TriLux, Perkin-Elmer).

### Grafting Experiment

The seedlings of wild type TL1, and the *cs1-cr1* mutant were grown under LD conditions with an intensity of 200 μmol·m^−2^·s^−1^ LED light at 25°C, for 7 days. The hypocotyl was surgically excised using a sterile blade for grafting. Subsequently, reciprocal grafting was performed with TL1 as a scion on *cs1-cr1* rootstock and vice versa. The graft union was securely sealed with Parafilm. After two days of postgrafting recovery in a humid and dark environment, the plants were returned to their original growth conditions.

### GUS Staining

The soybean cultivars NN1138-2 and TL1, along with the *cs1* and *cs1-cr1* mutants and the *CS1-OX-1* line, were subjected to transformation by Agrobacterium rhizogenes K599 bearing the *pDR5::GUS* plasmid to create transgenic hairy roots. The resulting hairy roots were then fixed in 90% (v/v) ice-cold acetone for a duration of 20 minutes and washed thrice with X-gluc-free staining solution, which comprised of 0.5 M potassium ferrocyanide, 0.5 M EDTA, 0.2 M phosphate buffer, 10 μl of 0.1% Triton X-100, 25 mg/ml, 0.5 mg/ml X-gluc, and 9.25 ml water. Subsequent to the rinsing, the roots were transferred to a 1.5 ml centrifuge tube containing X-gluc staining solution, ensuring complete submersion, and left undisturbed for a period of 1.5 hours at a temperature of 37℃ in a dark environment. The sample was subsequently decolorized using a decolorizing solution comprising 70% ethanol and 30% acetic acid. Following decolorization, the sample was inspected using a stereoscope and photographed.

### Low Blue Light Treatment

The seedlings were cultivated in SD conditions (12 h light/12 h dark), with a white light intensity of 550 μmol·m^−2^·s^−1^ at 25℃, for 12 days. Subsequently, the seedlings were subjected to a low blue light regime for 10 days to investigate the phenomenon of lodging. The low blue light condition consisted of a total light intensity of 350 μmol·m^−2^·s^−1^, with 16 μmol·m^−2^·s^−1^ of blue light. The control condition involved a normal white light intensity of 350 μmol·m^−2^·s^−1^, with 100 μmol·m^−2^·s^−1^ of blue light. The low blue light treatment was achieved by passing the white light through two layers of a yellow filter (no. 100, Lee Filters, CA), as previously described (Lyu et al., 2021). The quality and intensity of the light were measured using a HiPoint HR-350 spectrometer.

### Accession numbers

Sequence data from this article can be accessed in *phytozome*_13 libraries under accession numbers: *CS1* (*Glyma.19G186900*), *SGR6* (*AT2G36810*), expansin relative genes *Glyma.17G147600*, *Glyma.01G204800*, *Glyma.13G152200*, *Glyma.11G160000*, *Glyma.18G054000*, *Glyma.12G203600*, *Glyma.18G181700* and *Glyma.06G143300*, extensin relative genes (*Glyma.16G170000*, *Glyma.10G206900*, *Glyma.18G300800*, *Glyma.05G225300*, *Glyma.09G188600*, *Glyma.07G088300*, *Glyma.08G039400*, *Glyma.04G104600*, *Glyma.05G057500* and *Glyma.01G005600*), XTH relative genes (*Glyma.17G065200*, *Glyma.08G107300*, *Glyma.17G065300*, *Glyma.12G197600*, *Glyma.17G157400*, *Glyma.13G304400*, *Glyma.01G196800*, *Glyma.17G065100*, *Glyma.16G045000*, *Glyma.11G253900*, *Glyma.05G150400*, *Glyma.13G094900*, *Glyma.13G322500* and *Glyma.03G065500*), auxin transport genes (*Glyma.07G102500, Glyma.13G119000* and *Glyma.19G016700*), and auxin response genes (*Glyma.06G278400* and *Glyma.06G281900*). RNA-seq data from this article has been deposited in Genome Sequence Archive (GSA) under accession number PRJCA027194. The corresponding relationship between the names of the raw sequencing data and the materials in this paper are as follows: N38_2_A (NN1138-2-UH), N38_2_B (NN1138-2-MH), N38_2_C (NN1138-2-LH), *cs1*_A (*cs1*-UH), *cs1*_B (*cs1*-MH), *cs1*_C (*cs1*-LH). The soybean reference genome (*Glycine_max*.v2) was utilized in this study.

## SUPPLEMENTAL DATA

**Supplemental Figure S1.** Field phenotype of the *cs1* mutant.

**Supplemental Figure S2.** Diversity of agronomic traits between NN1138-2 and *cs1*.

**Supplementary Figure S3.** Hypocotyl Gravitropic response overtime in darkness.

**Supplementary Figure S4.** Gravitropic phenotypes of the *cs1* mutant and wild type NN1138-2 under light condition.

**Supplementary Figure S5.** Hypocotyl gravitropism patterns.

**Supplementary Figure S6.** Verification of mutation sites in isolated populations.

**Supplementary Figure S7**. Analysis of CS1 protein.

**Supplementary Figure S8.** Tissue-specific expression patterns of *CS1* in NN1138-2 and *cs1*.

**Supplementary Figure S9.** Comparison of *CS1* expression levels between the wild type (TL1) and over-expressing transgenic materials (*CS1-OX-1*, *CS1-OX-2*).

**Supplementary Figure S10.** Plant height comparison between the wild type (TL1) and transgenic lines (*CS1-OX-1*, *CS1-OX-2*).

**Supplementary Figure S11.** Amyloplast sedimentation in endodermal cells.

**Supplementary Figure S12.** The *CS1* gene regulates xylem and phloem development in the hypocotyl.

**Supplementary Figure S13.** The expression levels of the expansin, extensin and XTH relative gene families in NN1138-2 and *cs1*.

**Supplementary Figure S14.** The *cs1* mutation leads to a reduction in endoderm secondary cell wall thickness and in the content to cellulose and lignin in the hypocotyl.

**Supplementary Figure S15.** Volcanic plots of auxin biosynthesis, metabolism, transport and response related genes in MH and LH.

**Supplementary Figure S16.** Expression levels of auxin transport and auxin response gene families compared between NN1138-2 and *cs1*.

**Supplementary Figure S17.** Asymmetric distribution of IAA concentration in hypocotyls during Gravistimulation.

**Supplementary Figure S18.** Lodging rate assessment of grafted plants.

**Supplementary Figure S19.** Structural changes and hormone determination in the rootstock following grafting.

**Supplementary Figure S20.** Schematic diagram illustrating the regulatory role of *CS1* in preventing lodging.

## ACKNOWLEDGMENTS

We express our gratitude to Prof. Yonghong Wang, from the Institute of Genetics and Developmental Biology, Chinese Academy of Sciences, Beijing, China, for the invaluable assistance provided with the PAT assay. This work is supported by grants from the National Natural Science Foundation of China [32072091, 32171965], the Beijing Postdoctoral Research Foundation, Innovation Program of Chinese Academy of Agricultural Sciences, the Central Public-Interest Scientific Institution Basal Research Fund, the Core Technology Development for Breeding Program of Jiangsu Province (JBGS-2021-014), the Program of Collaborative Innovation Center for Modern Crop Production co-sponsored by Province and Ministry (CIC-MCP), and the National Key R & D Program of China (No. 2021YFD1201603).

## AUTHOR CONTRIBUTIONS

B.L., TJ.Z., T.Z. and Z.X. designed the research; Z.X., L.Z., K.K., and J.K. performed the experiments; Y.R., W.Z., J.L. and H.L. analyzed the data and participated in the experimental design; Z.X., TJ.Z. and B.L. wrote the paper; Y.R. provided experimental assistance; all the authors read and approved the manuscript.

---

**Supplementary Figure S1.**
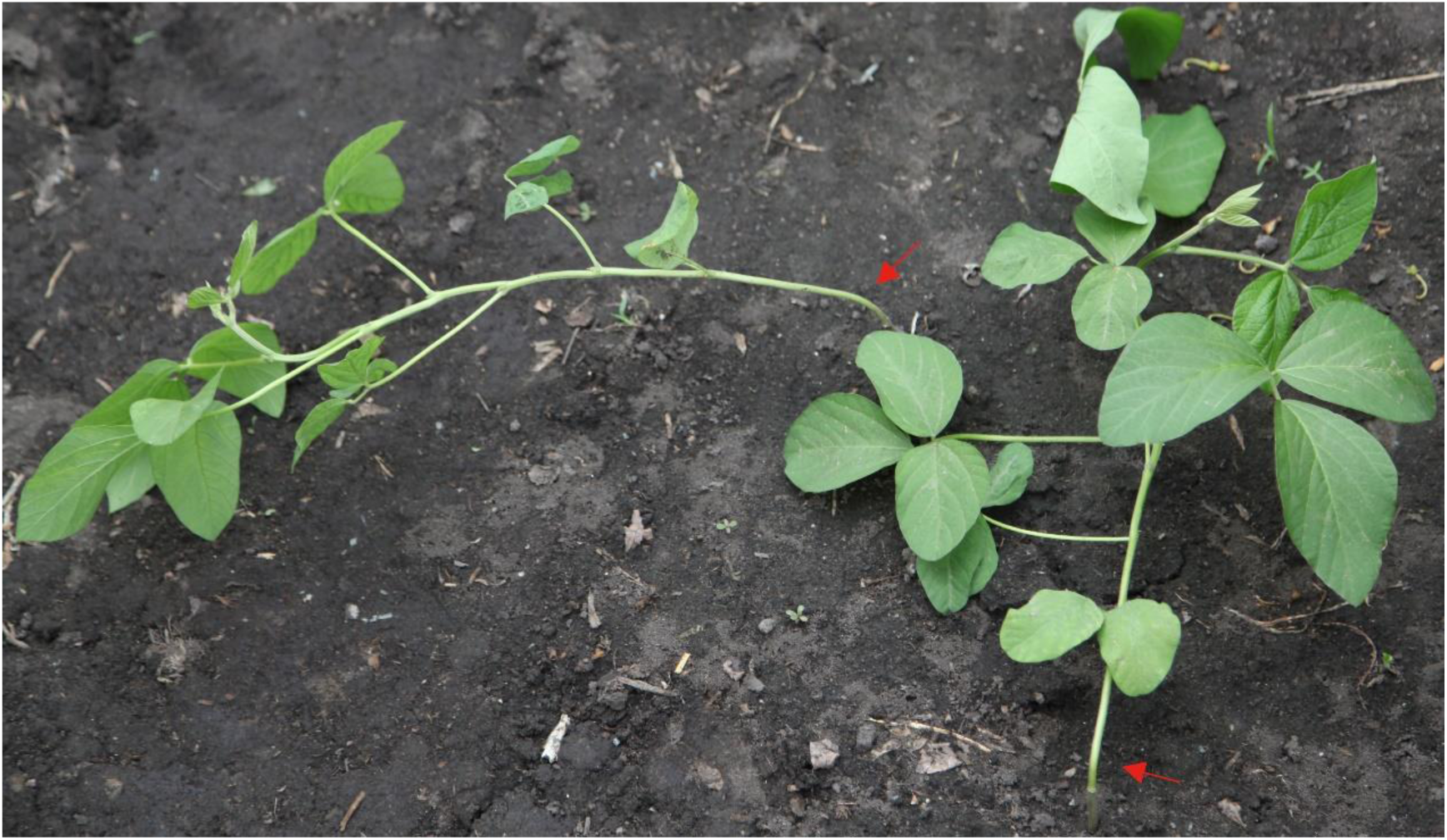
Field phenotype of the *cs1* mutant. The red arrow points to the specific lodging position.

**Supplementary Figure S2.**
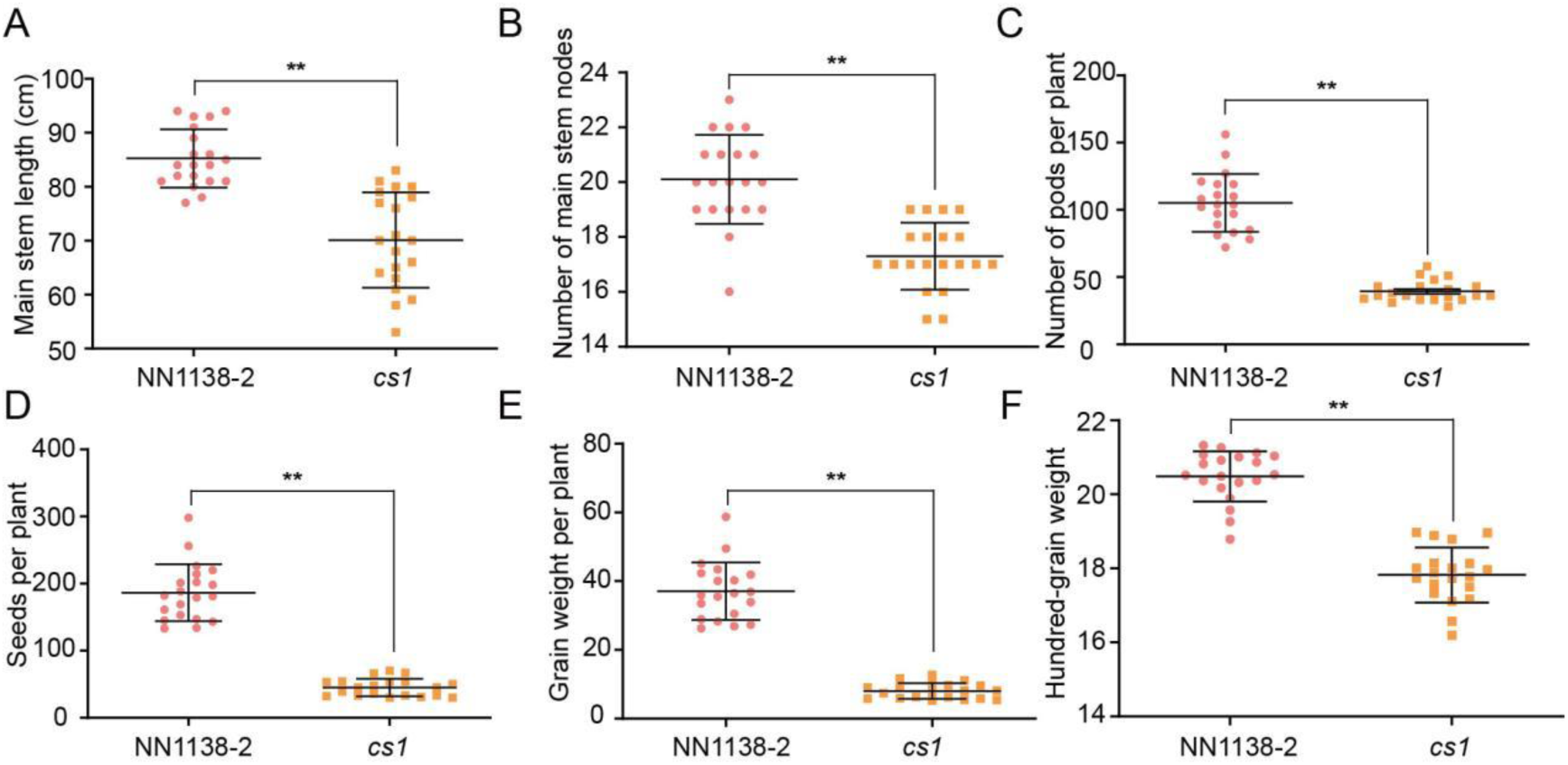
Diversity of agronomic traits between NN1138-2 and *cs1*. Comparison of main stem length (A), number of main stem nodes (B), number of pods per plant (C), seeds per plant (D), grain weight per plant (E) and hundred-grain weight (F) between NN1138-2 and *cs1*. The asterisks denote statistically significant differences from the wild type by a two-sided *t*-test (** P < 0.01). Values are means ± s.d. (*n* > 18).

**Supplementary Figure S3.**
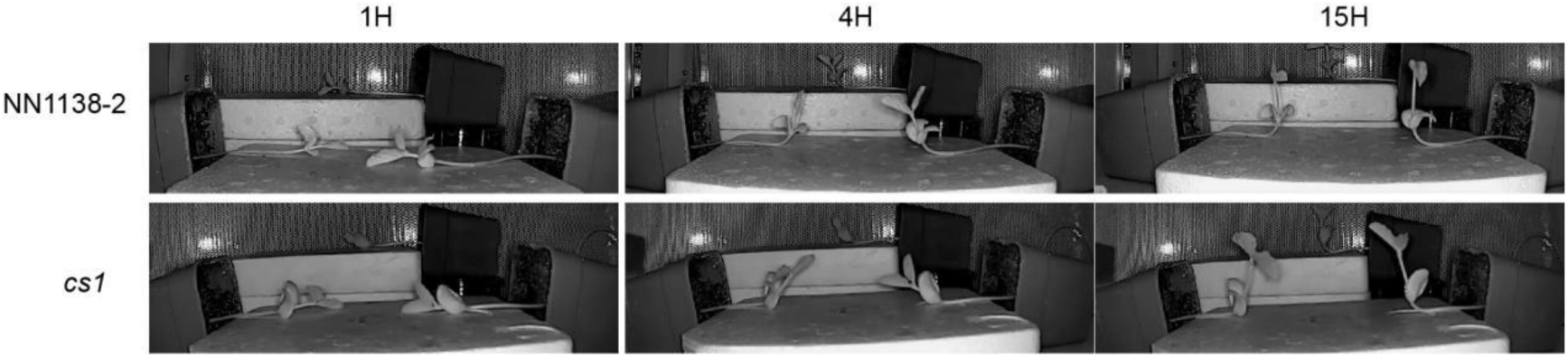
Hypocotyl Gravitropic response overtime in darkness. Seven-day-old seedlings were placed horizontally in darkness and the gravitropic response of hypocotyl were recorded at intervals of 1 hour, 4 hours and 15 hours.

**Supplementary Figure S4.**
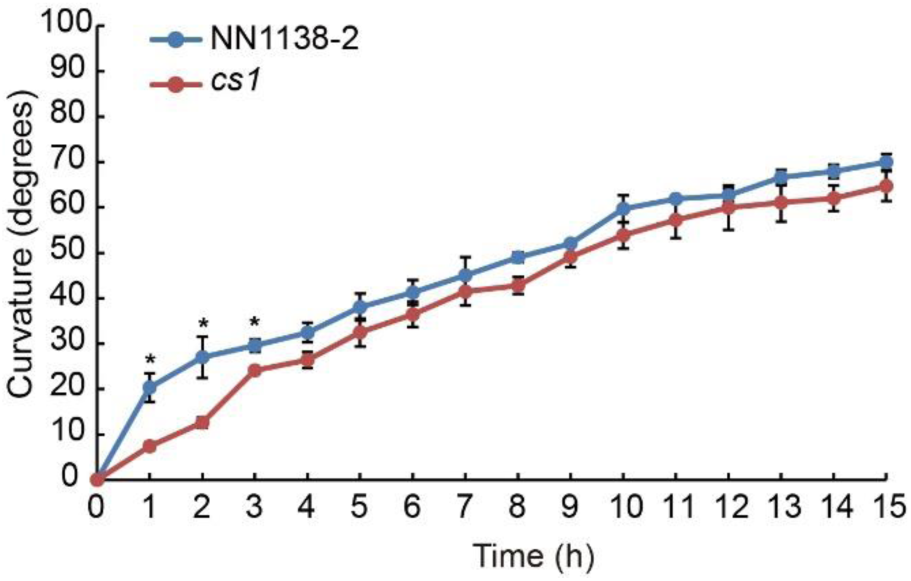
Gravitropic phenotypes of the *cs1* mutant and wild type NN1138-2 under light condition. Values are means ± SD (*n* = 3). The asterisks indicate statistically significant difference from the wild type as determined by a two-sided *t*-test (* P < 0.05).

**Supplementary Figure S5.**
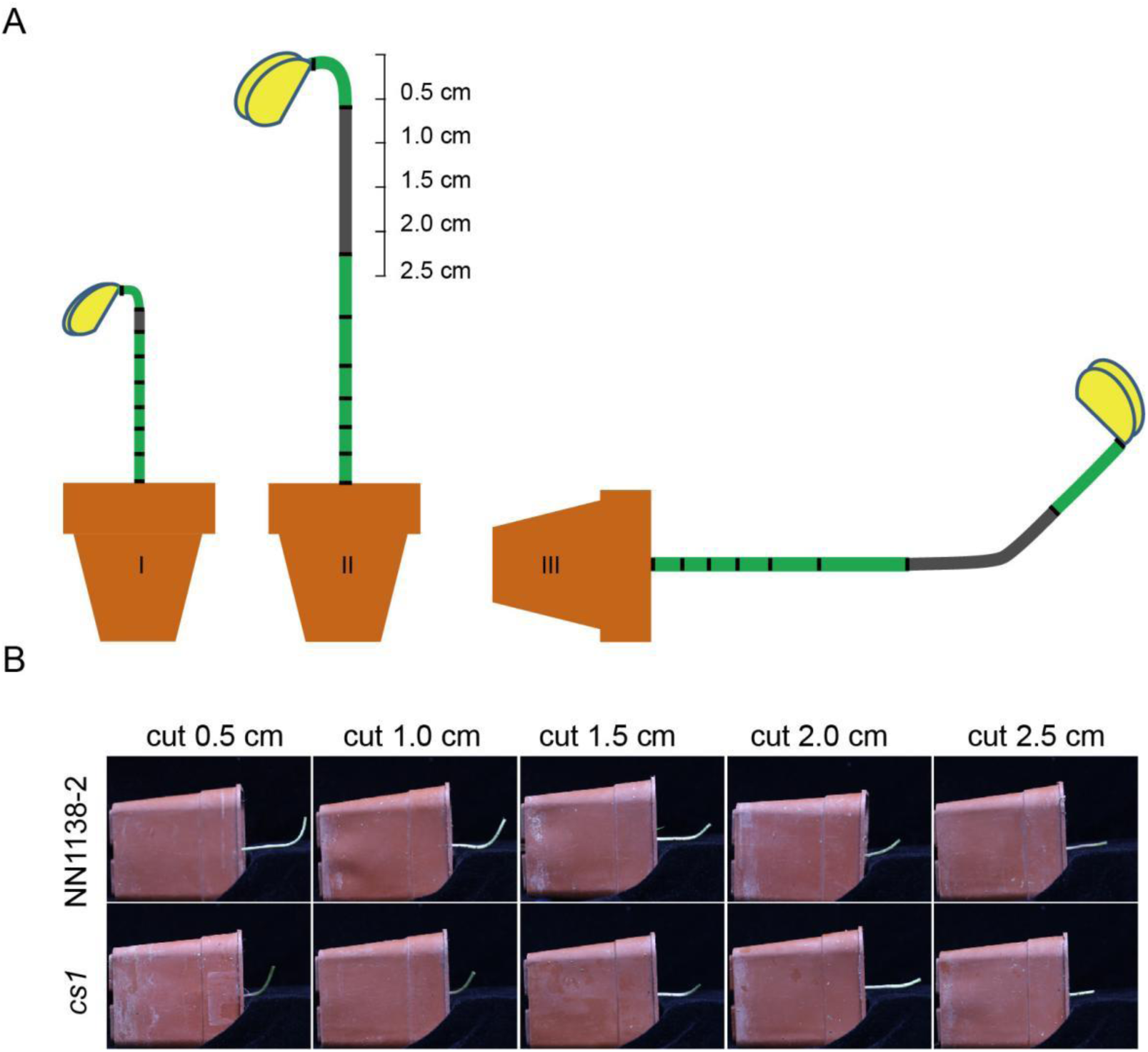
Hypocotyl gravitropism patterns. **A)** Hypocotyl gravitropism pattern diagram. I: Plants grown for 3.5 days were marked at intervals of 0.5 cm from the growth point. II: The distances between each mark were measured after 24 hours, and the hypocotyls were sectioned into 0.5 cm increments from the growth point. III: The sectioned plants were placed horizontally in darkness. After 10 hours, the bending at the marked portions was observed. The bend is indicated in gray. **B)** Bending photographs of the sectioned hypocotyls of indicated lines.

**Supplementary Figure S6.**
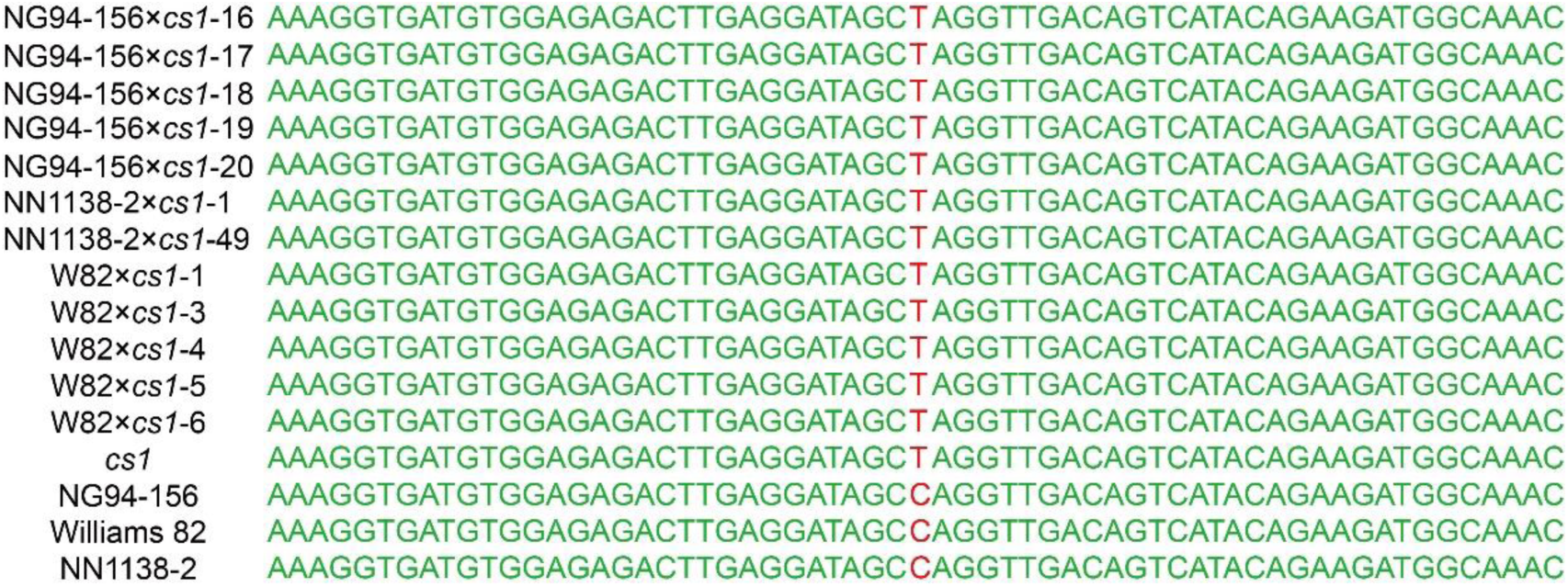
Verification of mutation sites in isolated populations. The isolated population of lodging plants showed consistency with the parental mutant mutation sites. Red indicates the SNP site, with base C representing the wild type and T representing the mutant plants respectively.

**Supplementary Figure S7.**
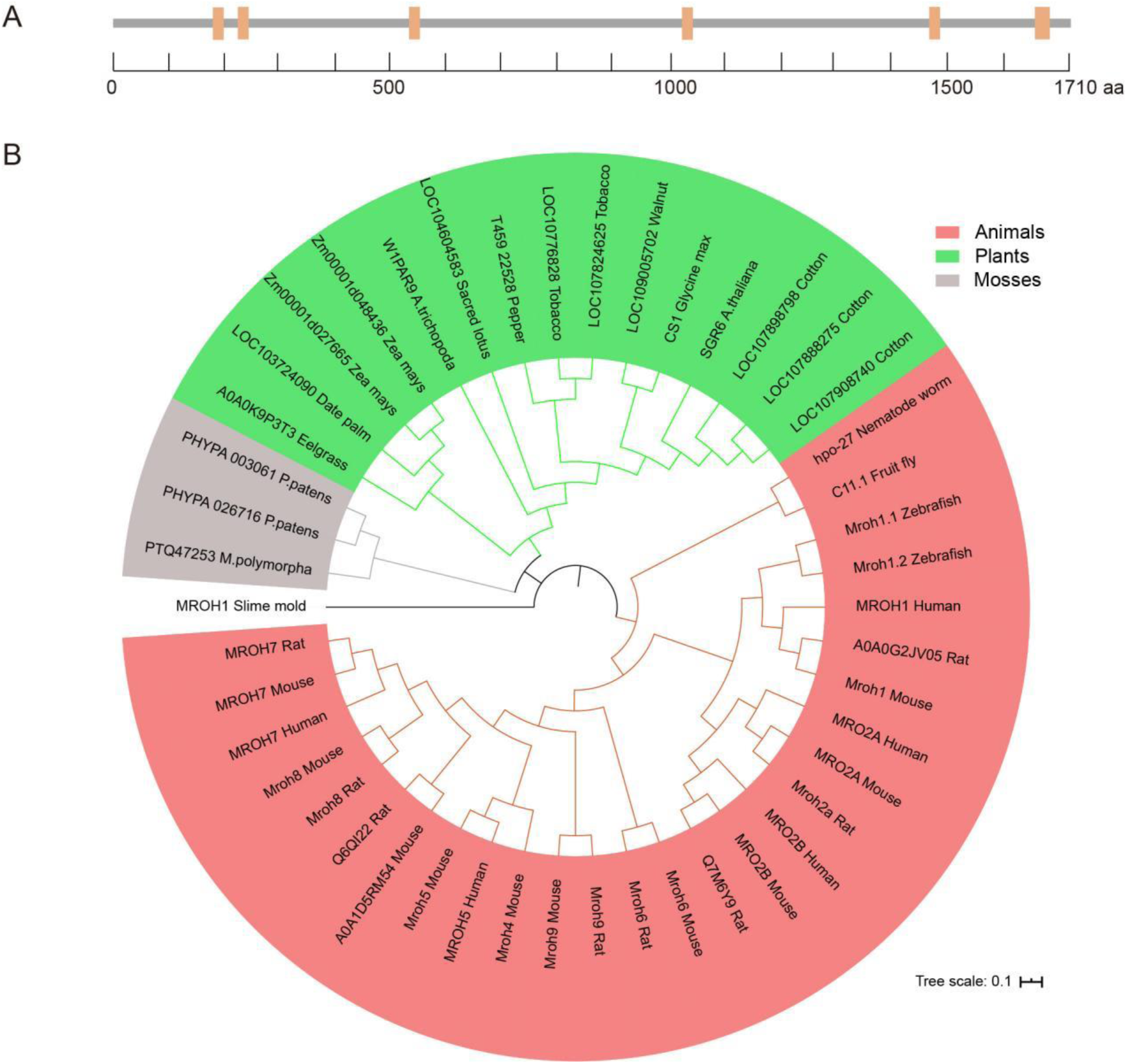
Analysis of CS1 protein. **A)** CS1 protein sequence length pattern in wild type. An orange box represents a HEAT repeat within the protein sequencing. **B)** Phylogenetic tree of CS1 across animals, plants, and mosses.

**Supplementary Figure S8.**
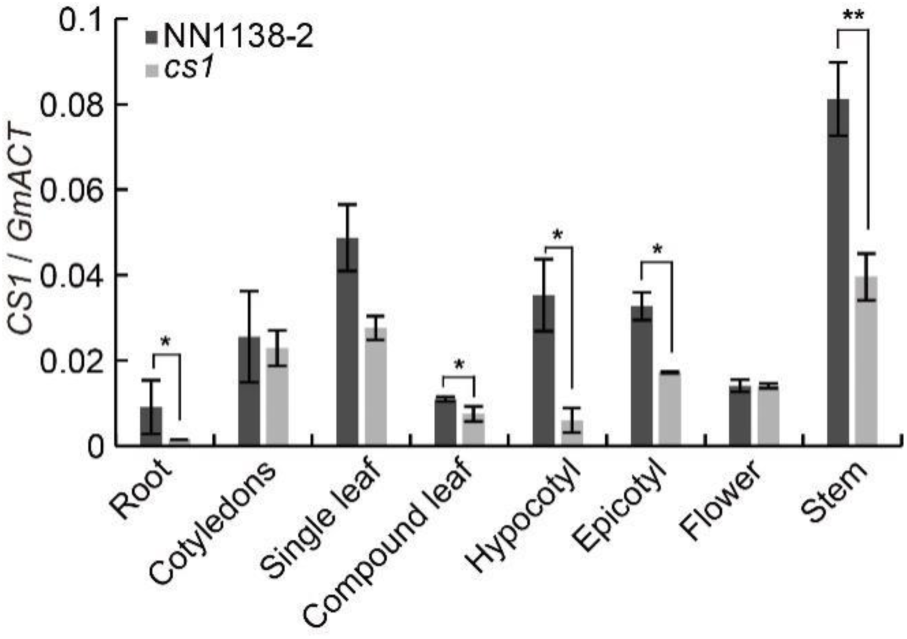
Tissue-specific expression patterns of *CS1* in NN1138-2 and *cs1*. *GmActin11* was used as a control in the real-time PCR analyses. The asterisks indicate statistically significant difference from the wild type by two-sided *t*-test (* *P* < 0.05; ** *P* < 0.01).

**Supplementary Figure S9.**
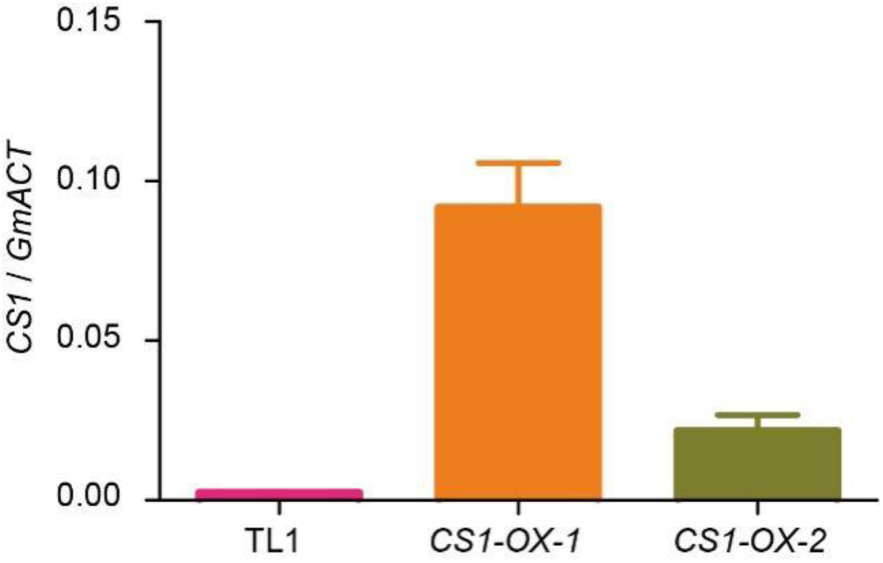
Comparison of *CS1* expression levels between the wild type (TL1) and over-expressing transgenic materials (*CS1-OX-1*, *CS1-OX-2*). *GmActin11* was used as a control gene in the real-time PCR analyses. Values are means ± s.d. (*n* = 3).

**Supplementary Figure S10.**
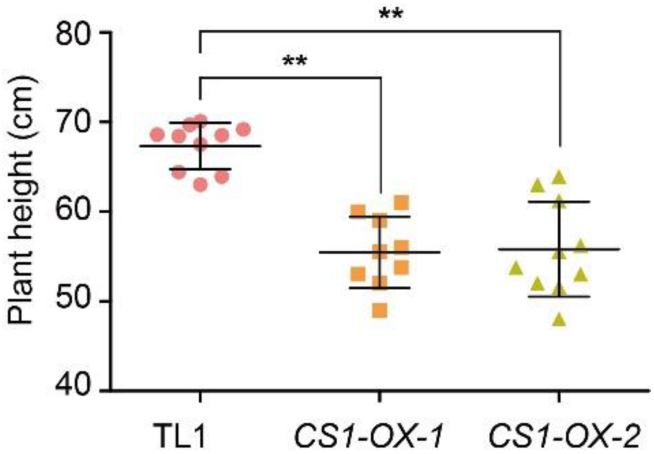
Plant height comparison between the wild type (TL1) and transgenic lines (*CS1-OX-1*, *CS1-OX-2*). Values are means ± s.d. (*n* > 9). The asterisks indicate statistically significant differences from the wild type by two-sided *t*-test (** P < 0.01).

**Supplementary Figure S11.**
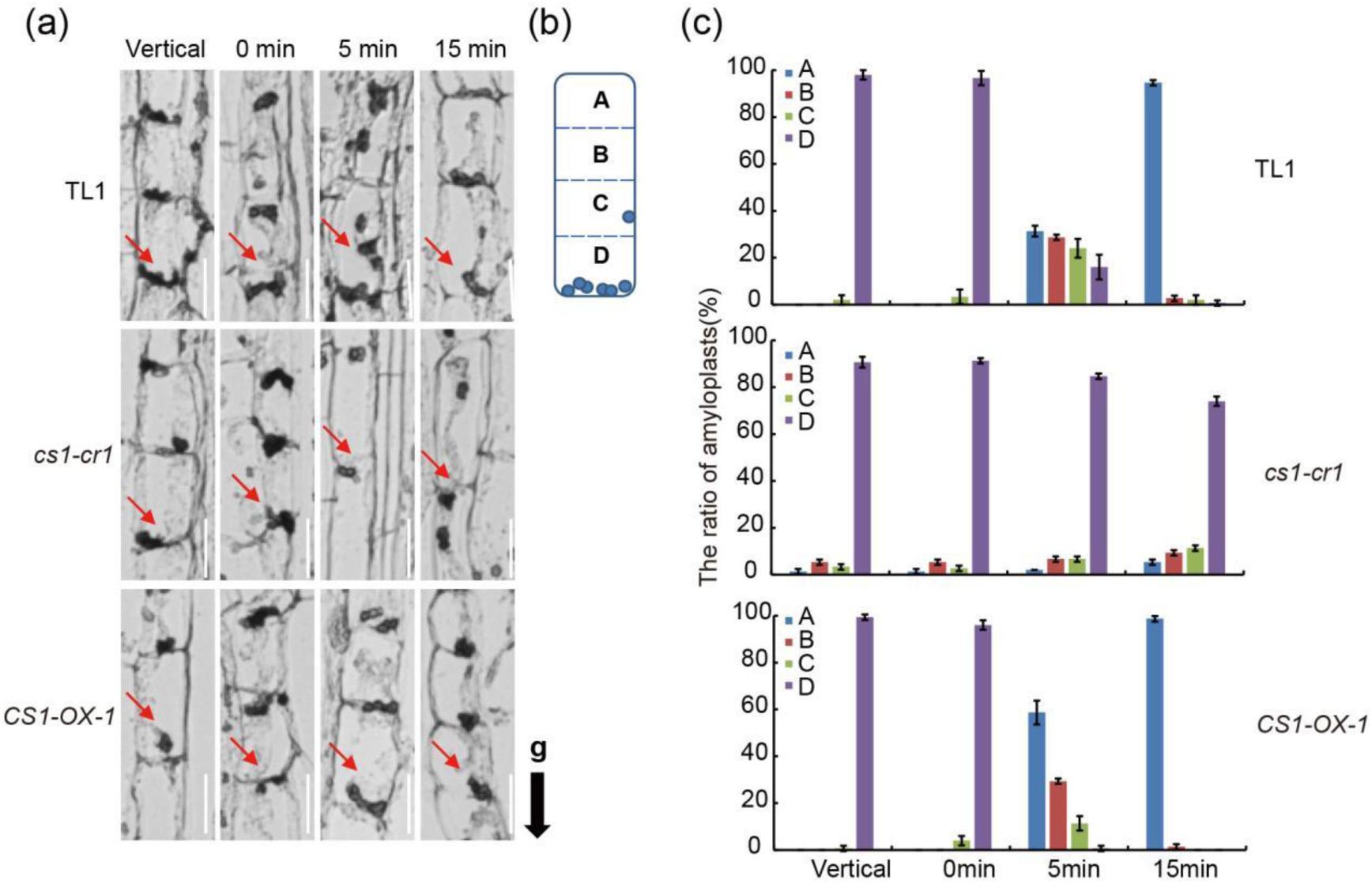
Amyloplast sedimentation in endodermal cells. **A)** Longitudinal section images show the distribution of amyloplasts in endodermal cells (similar to Figure 1). Seven-day-old vertically growing plants were inverted for indicated durations (0 to 15 min). Hypocotyl fragments fragments (1-2 cm below the cotyledon node) were collected and fixed, with the gravity direction maintained at the indicated time point. Red arrows indicate the locations of the amyloplasts, and the black arrow indicates the direction of gravity (g). Scale bars = 5 um. **B-D)** The ratio of amyloplasts in each block for TL1 (B), *cs1-cr1* (C) and *CS1-OX-1*(D). Data are means ± s.d. (*n* = 50).

**Supplementary Figure S12.**
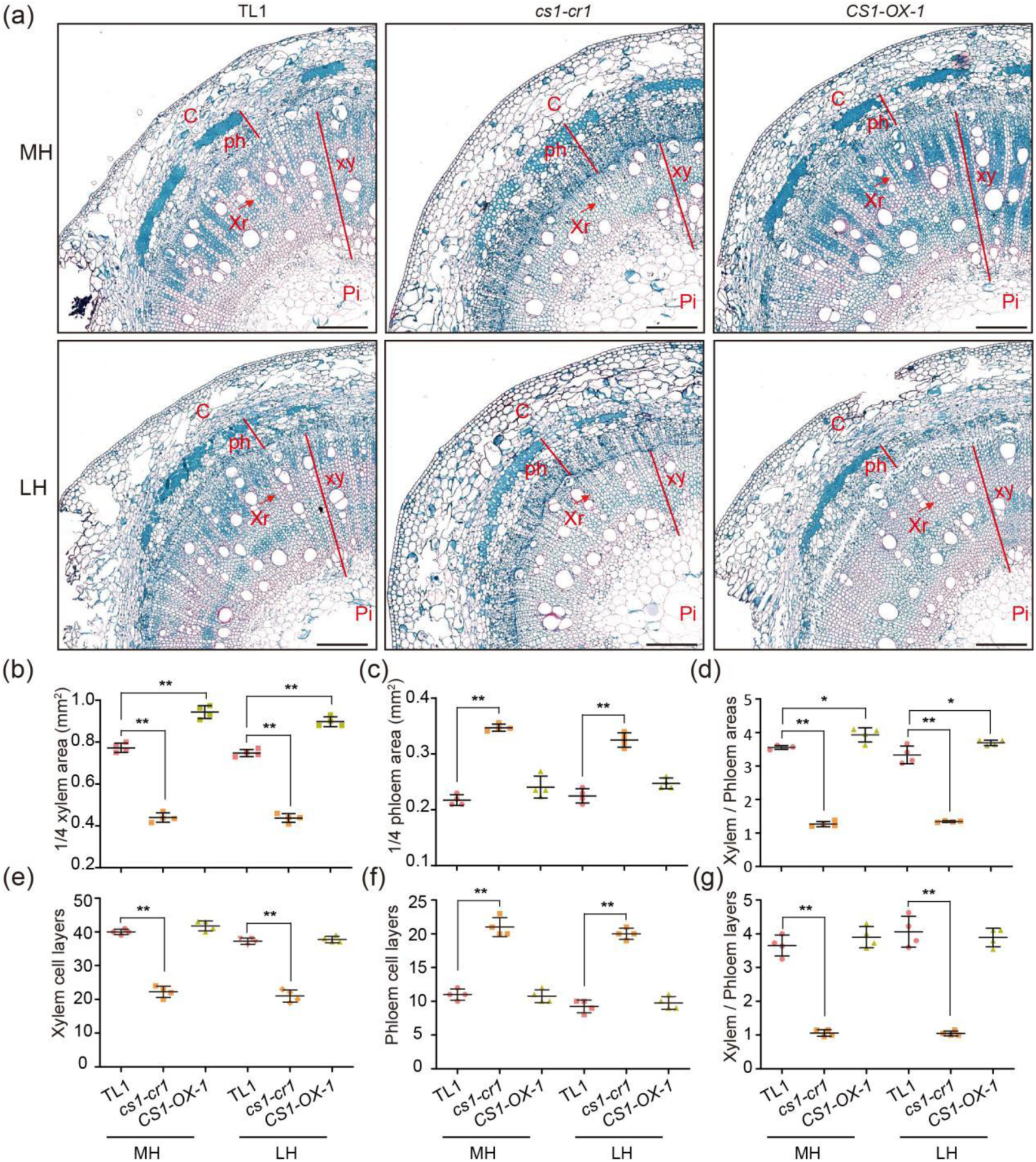
The *CS1* gene regulates xylem and phloem development in the hypocotyl. **A)** Quadrant cross-sectional images show the cellular layer structures in the middle and lower regions of the hypocotyl. Plants of the indicated genotypes were grown under LD conditions for 7 days. C, cortex; ph, phloem; xr, xylem rays; xy, xylem; pi, pith. Red arrows indicate xylem rays. Scale bar = 250 um. **B-G)** Scatter plots of xylem areas (B), phloem areas (C), xylem areas / phloem areas (D), xylem cell lays (E), phloem cell lays (F), and xylem cell lays / phloem cell lays (G) of indicated lines as in (A). Data are means ± s.d. (*n* = 4). The asterisks indicate statistically significant difference from the wild type by two-sided *t*-test (* *P* < 0.05; ** *P* < 0.01).

**Supplementary Figure S13.**
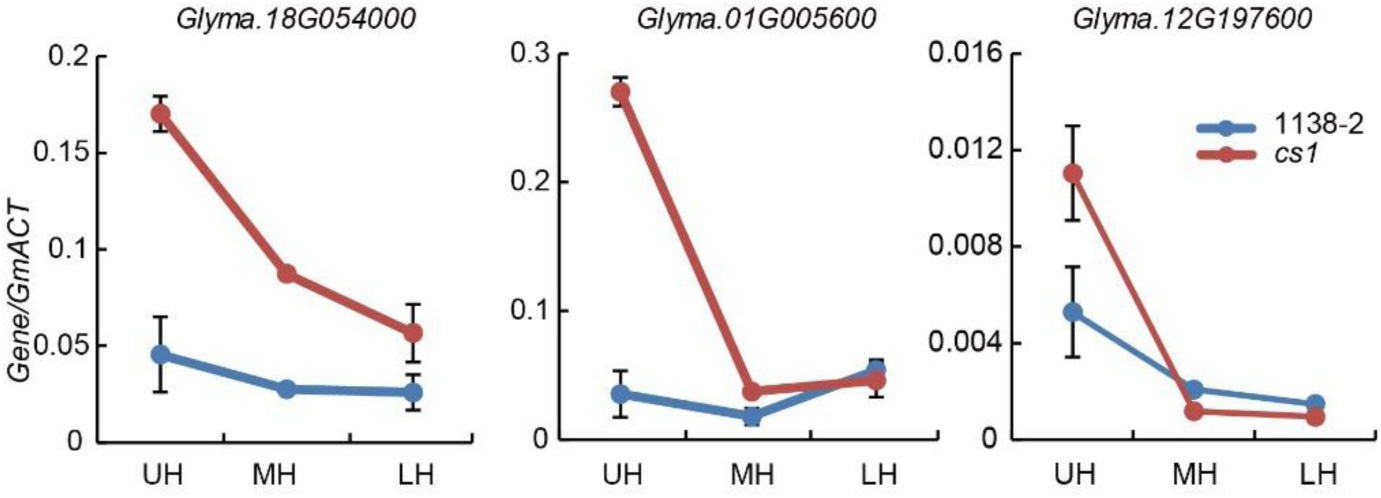
The expression levels of the expansin, extensin and XTH relative gene families in NN1138-2 and *cs1*. *GmActin11* was used as a control in the real-time PCR analyses. Values are means ± s.d. (*n* = 3).

**Supplementary Figure S14.**
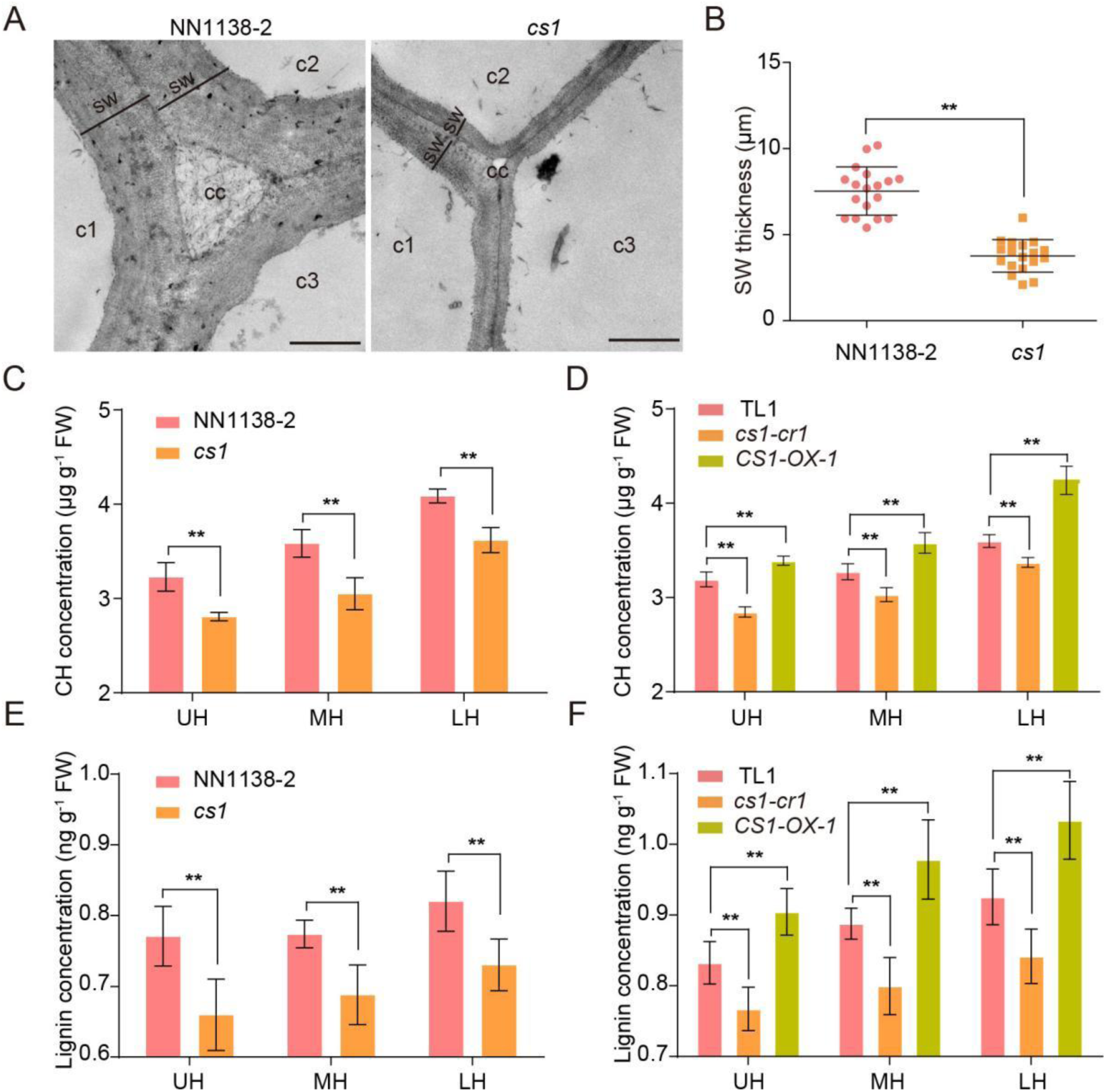
The *cs1* mutation leads to a reduction in endoderm secondary cell wall thickness and in the content to cellulose and lignin in the hypocotyl. **A)** Transmission electron micrographs (TEM) of the upper hypocotyl endodermal cell walls from indicated NIL lines. Scale bar = 6 µm. C1∼3: cell 1∼3, sw: secondary cell wall, cc: cell corners, va: vacuole, ap: amyloplast. **B)** The Upper hypocotyl endodermal secondary cell wall thickness (statistical results from a). Values are means ± s.d. (*n* = 18). **C-D)** Cellulose and hemicellulose (CH) concentration in hypocotyl of indicated genotypes. FW means fresh weight. **E-F)** Lignin concentration in hypocotyl of indicated genotypes. Values are means ± s.d. (*n* ≥ 4). The asterisks indicate statistically significant difference from the wild type by two-sided *t*-test (* *P* < 0.05; ** *P* < 0.01).

**Supplementary Figure S15.**
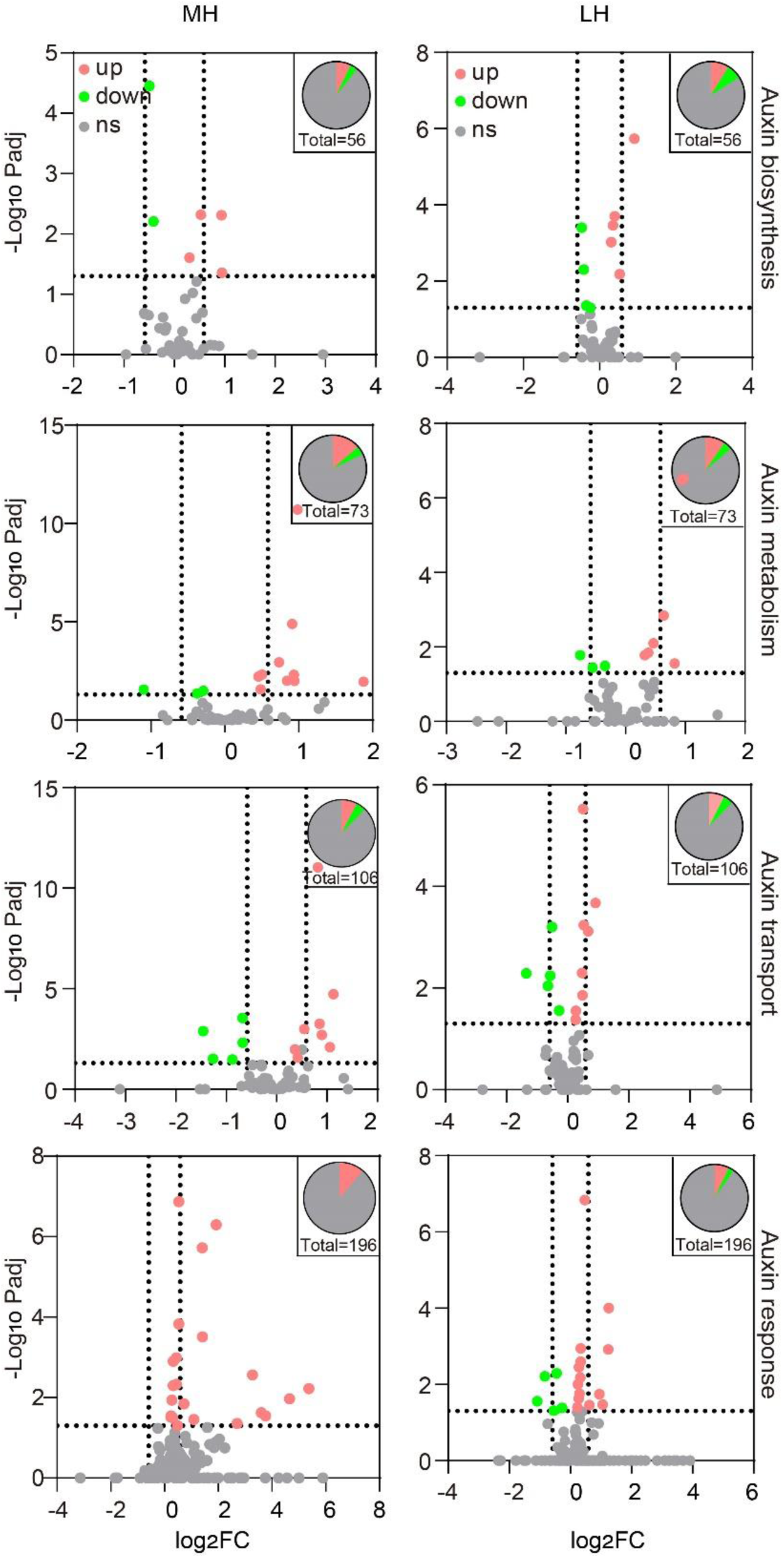
Volcanic plots of auxin biosynthesis, metabolism, transport and response related genes in MH and LH.

**Supplementary Figure S16.**
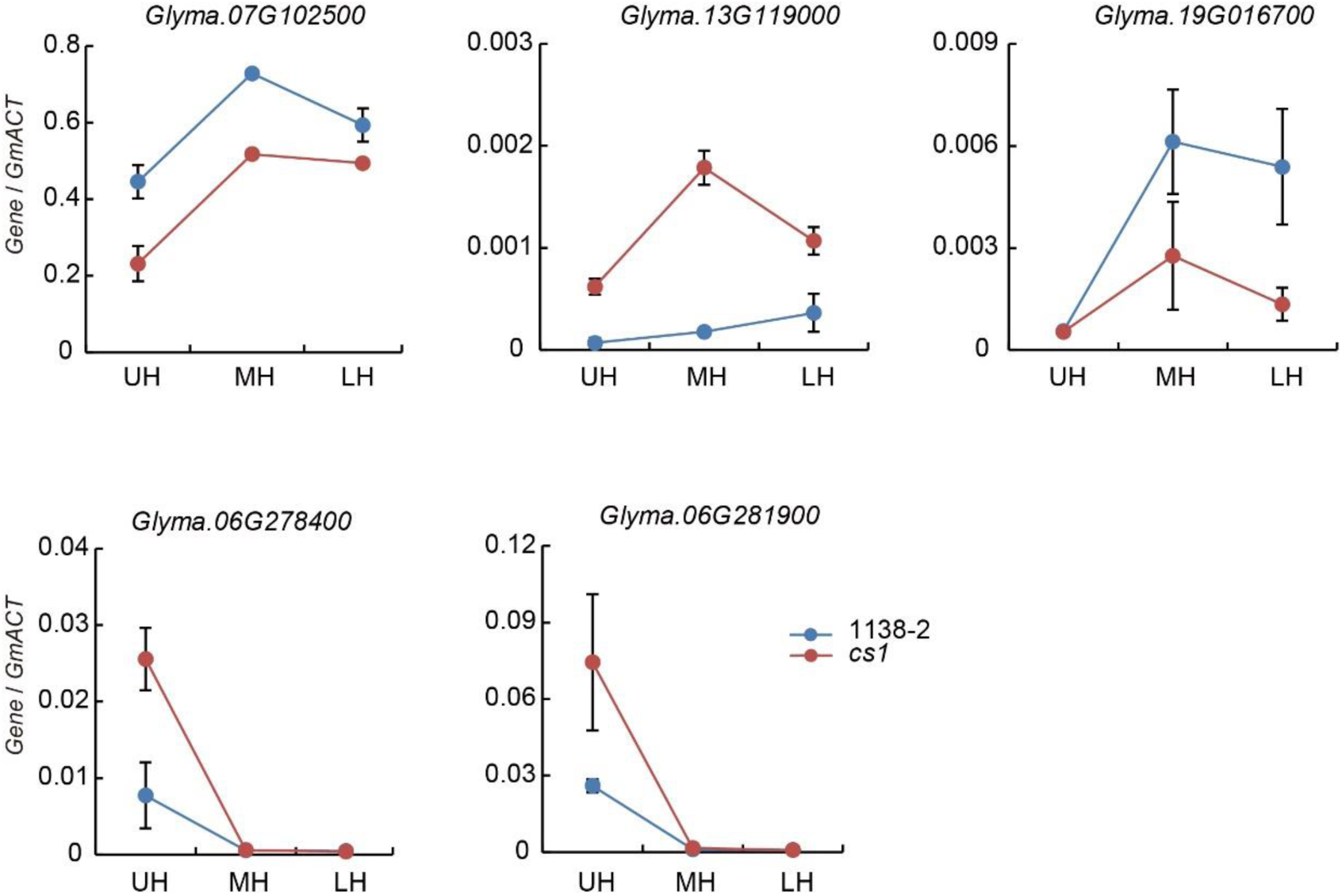
Expression levels of auxin transport and auxin response gene families compared between NN1138-2 and *cs1*. *GmActin11* was used as a control in the real-time PCR analyses. Values are means ± s.d. (*n* = 3).

**Supplementary Figure S17.**
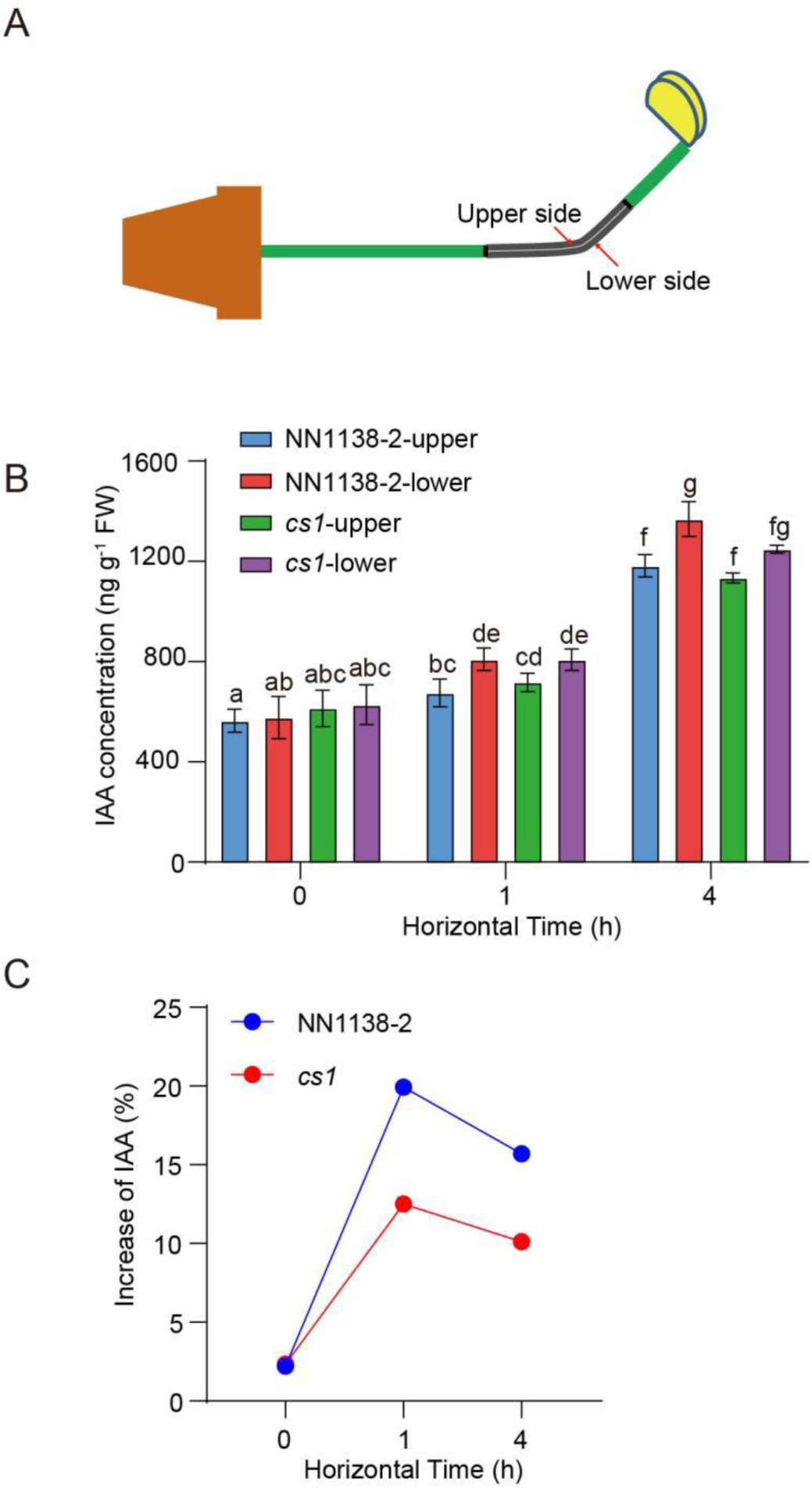
Asymmetric distribution of IAA concentration in hypocotyls during Gravistimulation. **A)** Seven-day-old soybean seedlings were rotated 90° from the vertical to horizontal position and incubated for 0, 1, and 4 h, respectively. The gray area represents the site that was harvested at each time point. Hypocotyls were divided into their lower and upper sides along the midline (indicated by a white line). **B)** IAA concentration on the upper and lower sides of NN1138-2 and *cs1* after gravistimulation. FW means fresh weight. Values are means ± s.d. (*n* ≥ 4). **C)** The increase in IAA after gravistimulation is shown as percentage change, calculated as [(lower-upper)/upper]*100%.

**Supplementary Figure S18.**
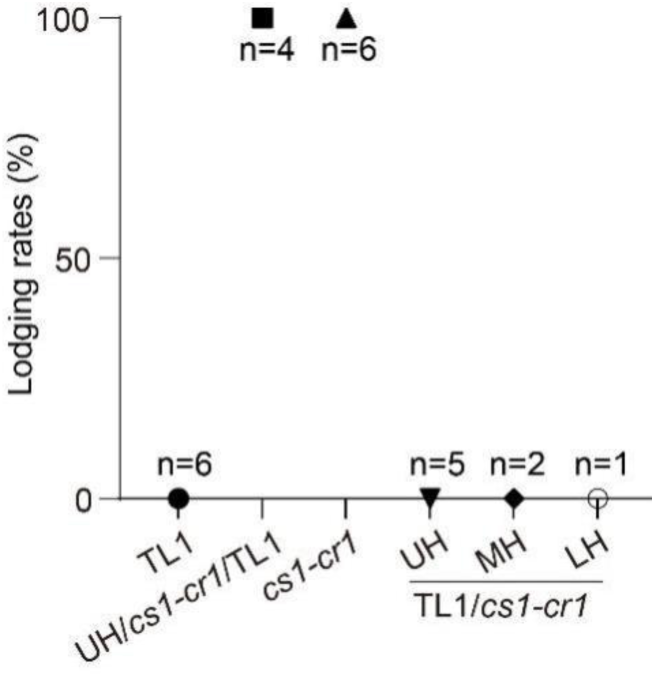
Lodging rate assessment of grafted plants. UH, MH and LH are the grafting sites, n is the number of grafted plants.

**Supplementary Figure S19.**
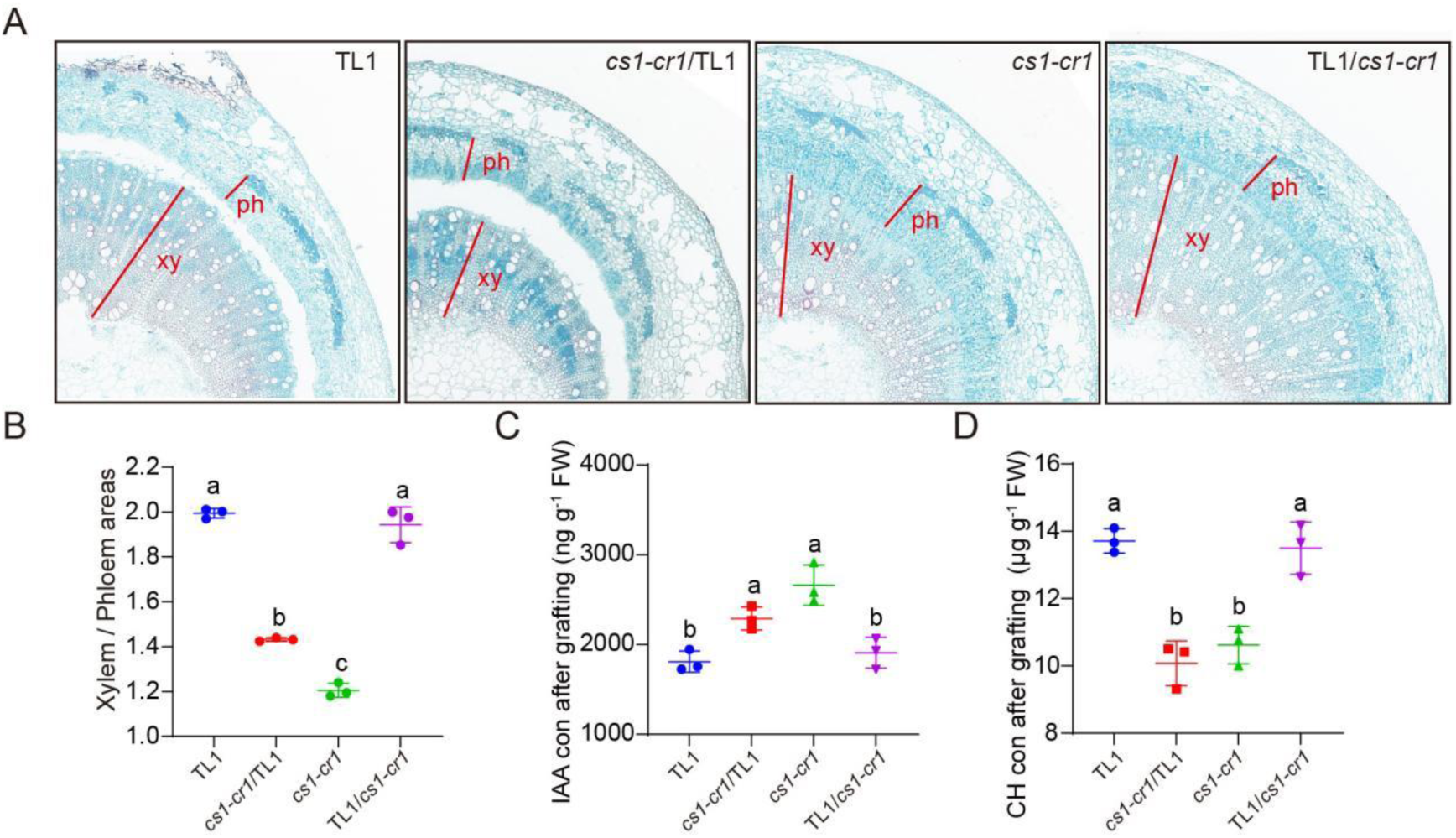
Structural changes and hormone determination in the rootstock following grafting. **A)** Quadrant cross-sectional images show the cell layer structures within the hypocotyl rootstock. Ph, phloem; xy, xylem. **B)** Scatter plots represent the ratio of xylem areas to phloem areas. **C)** Auxin concentration measurements in hypocotyl rootstock of indicated lines. **D)** Cellulose and hemicellulose (CH) concentration in hypocotyl rootstock of indicated lines. *cs1-cr1*/TL1: *cs1-cr1* is scion, TL1 is rootstock; TL1/ *cs1-cr1*:TL1 is scion, *cs1-cr1* is rootstock. The letters above the bars indicate significant differences (*P* < 0.05) as determined by one-way ANOVA following by Tukey’s multiple comparisons test.

**Supplementary Figure S20.**
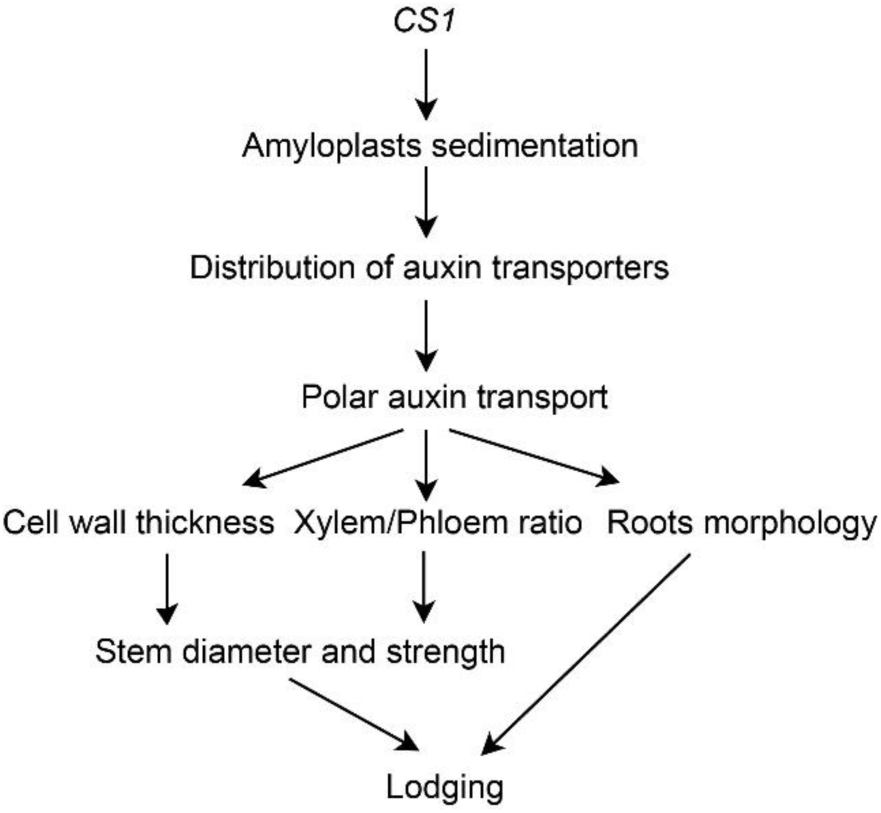
Schematic diagram illustrating the regulatory role of *CS1* in preventing lodging. Note: The arrows indicates the direction and correlation between the various factors influencing lodging resitance.

**Supplementary Table 1:** Goodness-of-fit test for segregation ratios of erect to creeping trait in four crosses between the normal parents and the mutant *cs1*.

| Population | No. of WT plants | No. of MT plants | Expected ratio | $\chi^2$ | P |
| --- | --- | --- | --- | --- | --- |
| Williams 82 × *cs1* | 1251 | 212 | 3:1 | 85.62 | 0.00 |
| NG94-156 × *cs1* | 209 | 45 | 3:1 | 6.80 | 0.01 |
| KF1 × *cs1* | 485 | 98 | 3:1 | 20.42 | 0.00 |
| NN1138-2 × *cs1* | 548 | 116 | 3:1 | 19.68 | 0.00 |
| | No. of segregating lines | No. of non-segregating lines | Expected ratio | $\chi^2$ | P |
| NN1138-2 × *cs1* F<sub>2:3</sub> | 34 | 17 | 2:1 | 0.02 | 0.88 |
| KF1 × *cs1* F<sub>2:3</sub> | 29 | 15 | 2:1 | 0.01 | 0.94 |

**Supplementary Table 2:**
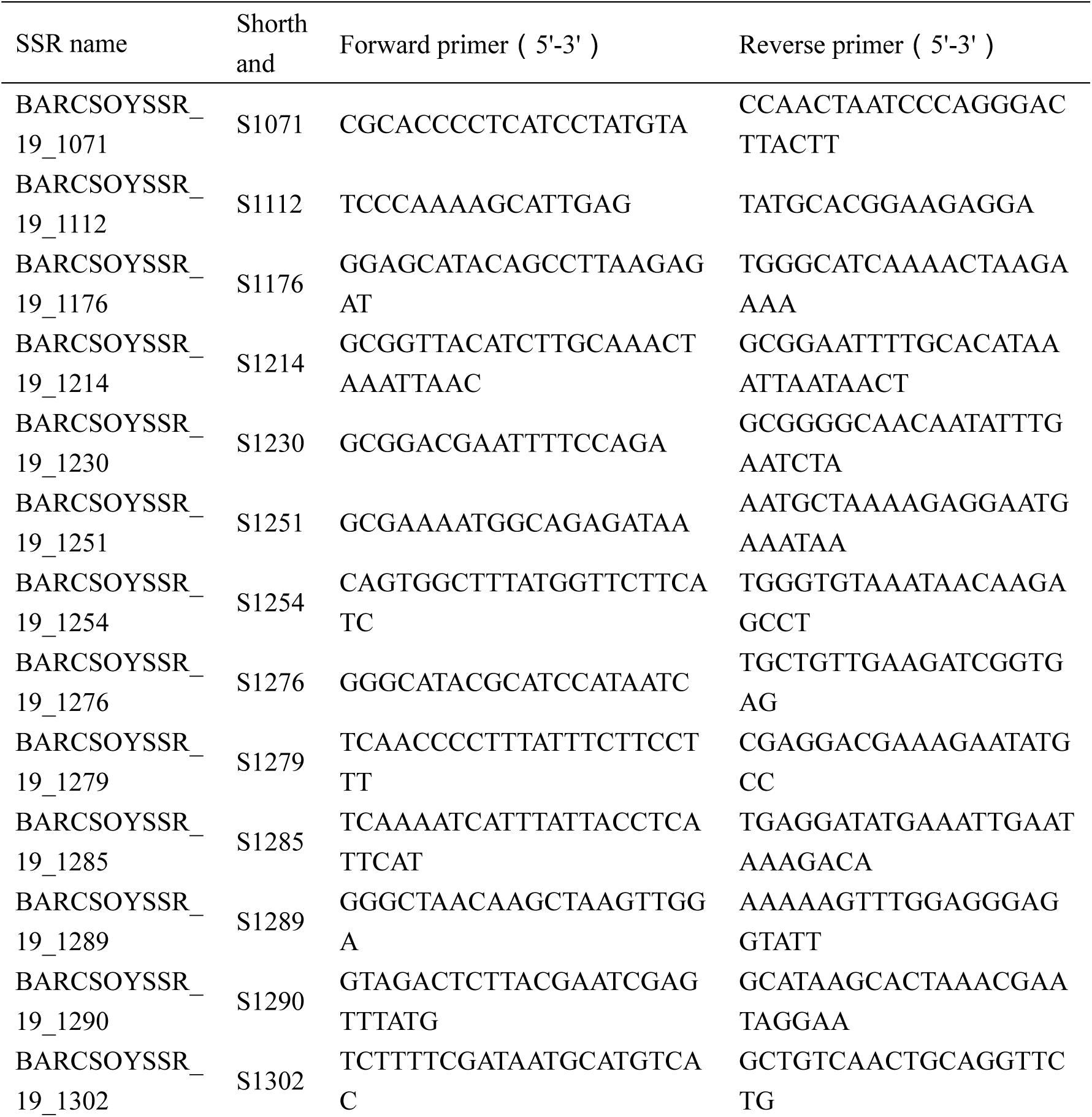
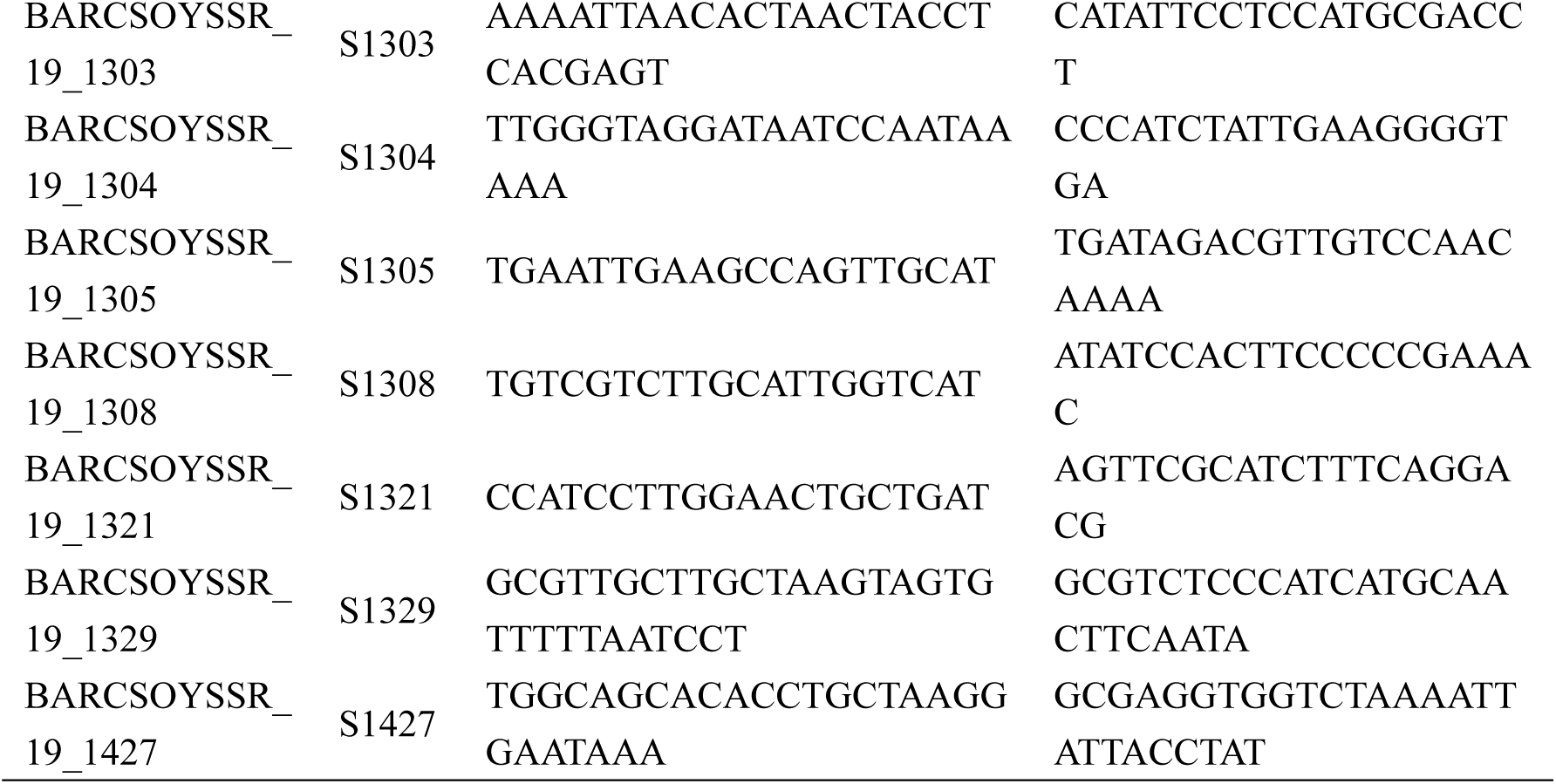
Names and corresponding primer sequences of SSR markers for mapping *cs1* gene.

**Supplementary Table 3:**
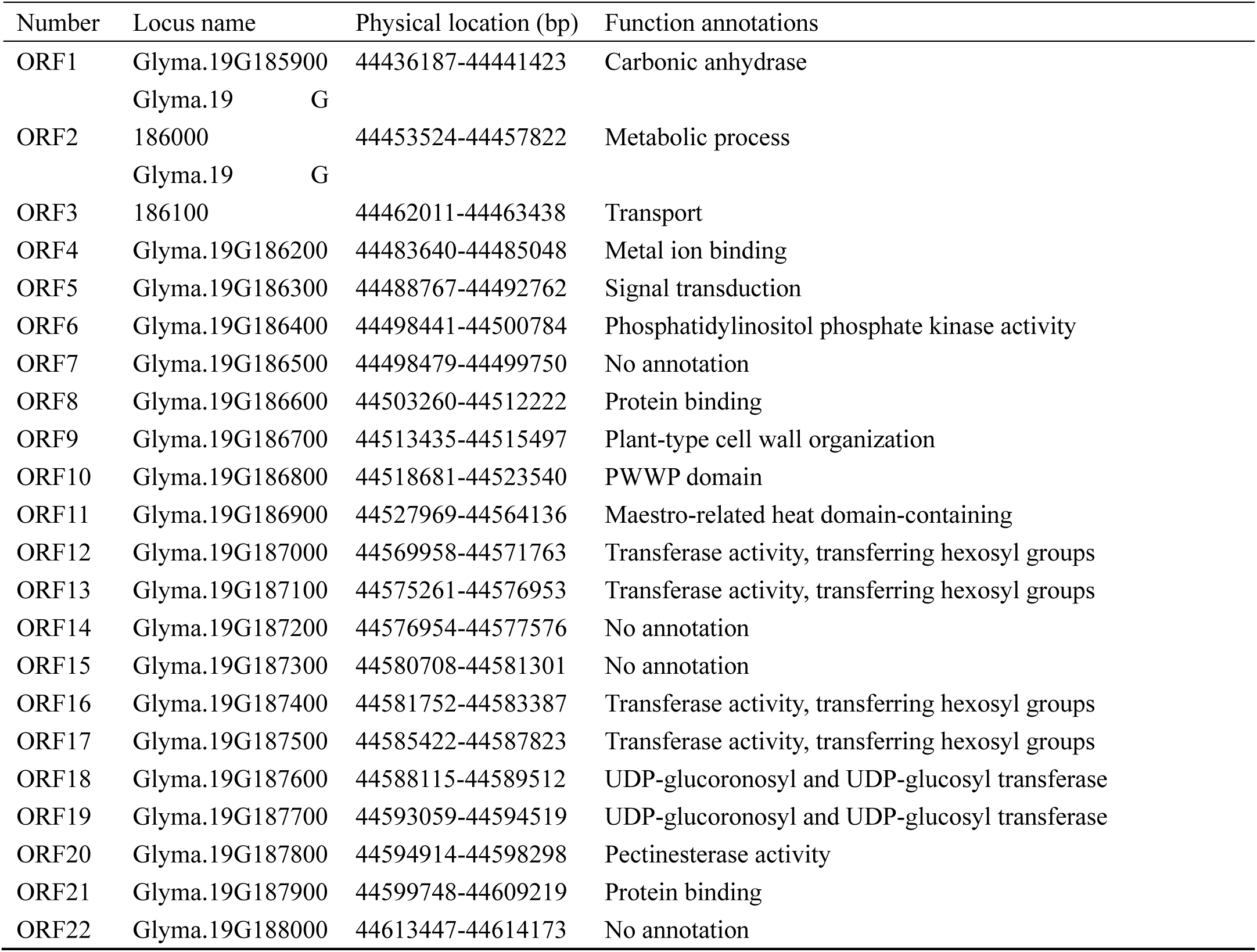
Candidate genes and corresponding functions in the target mapping region of *cs1*.

